# PP2A-B56 regulates Mid1 protein levels for proper cytokinesis in fission yeast

**DOI:** 10.1101/2024.06.28.601230

**Authors:** Madeline L. Chrupcala, James B. Moseley

**Affiliations:** Department of Biochemistry and Cell Biology, The Geisel School of Medicine at Dartmouth, Hanover NH

## Abstract

Protein phosphorylation regulates many steps in the cell division process including cytokinesis. In fission yeast cells, the anillin-like protein Mid1 sets the cell division plane and is regulated by phosphorylation. Multiple protein kinases act on Mid1, but no protein phosphatases have been shown to regulate Mid1. Here, we discovered that the conserved protein phosphatase PP2A-B56 is required for proper cytokinesis by promoting Mid1 protein levels. We find that *par1Δ* cells lacking the primary B56 subunit divide asymmetrically due to the assembly of misplaced cytokinetic rings that slide towards cell tips. These *par1Δ* mutants have reduced whole-cell levels of Mid1 protein, leading to reduced Mid1 at the cytokinetic ring. Restoring proper Mid1 expression suppresses *par1Δ* cytokinesis defects. This work identifies a new PP2A-B56 pathway regulating cytokinesis through Mid1, with implications for control of cytokinesis in other organisms.

## INTRODUCTION

Cytokinesis physically separates a mother cell into two functional daughter cells, thereby completing the cell cycle. This process must be tightly regulated in both time and space to ensure the viability of each daughter cell. Given this need, cells have evolved robust mechanisms to promote efficient cytokinesis, and these mechanisms are conserved across eukaryotes (Pollard & O’Shaughnessy, 2019). Cytokinesis in both fungi and animal cells occurs in a series of defined events (Bhavsar-Jog & Bi, 2017; Cheffings et al., 2016; D’Avino et al., 2015; Green et al., 2012; Mangione & Gould, 2019; Pollard & Wu, 2010). The cell initially selects the division plane. Next, cells assemble a contractile actomyosin ring (CAR) at this site. Finally, the CAR constricts and subsequently disassembles. These events culminate in the physical separation of the cell. While much of the CAR machinery has been identified and is evolutionarily conserved from yeast to humans, many questions remain open regarding how these proteins assemble and constrict the CAR in a regulated manner.

Protein phosphorylation is an important mechanism that regulates CAR proteins in both time and space (Almonacid et al., 2009, 2011; Bähler, Steever, et al., 1998; Barr et al., 2004; Burkard et al., 2009; Kim et al., 2017; Magliozzi et al., 2020; Rezig et al., 2023; Willet et al., 2019a; Wolfe et al., 2009). Many protein kinases have been shown to regulate both temporal and spatial steps of cytokinesis in eukaryotes (Barr et al., 2004; Bohnert & Gould, 2011; Brennan et al., 2007; Burkard et al., 2009; Carmena et al., 2012; Magliozzi & Moseley, 2021; Santamaria et al., 2007; Sechi et al., 2022; Su et al., 2011; van der Waal et al., 2012; Wolfe et al., 2009). The level of phosphorylation on a protein is established by the balanced activities of protein kinases and protein phosphatases. Some protein phosphatases have been connected with regulation of cytokinesis (Barr et al., 2004; Burgess et al., 2010; Cundell et al., 2016; García-Blanco et al., 2019; Holder et al., 2019). For example, budding yeast PP2A-Cdc55 has been identified as a regulator of CR contraction and septum assembly during cytokinesis (Moyano-Rodríguez et al., 2022). In addition, Calcineurin/PP2B and Clp1 localize to the CAR in fission yeast and regulate cytokinesis (Martín-García et al., 2018; Mishra et al., 2004; Snider et al., 2020; Trautmann et al., 2001; Yoshida et al., 1994). However, protein phosphatases remain far less investigated in cytokinesis compared to protein kinases.

The fission yeast *Schizosaccharomyces pombe* has proven to be a strong model organism for dissecting the complex mechanisms underlying cytokinesis. Following mitosis, these rod-shaped cells divide precisely in the cell middle to generate two equally sized daughter cells. Fission yeast cytokinesis follows a reproducible pattern of three steps. First, the CAR is formed by a series of multiprotein “nodes” that bind to the plasma membrane (Almonacid et al., 2009, 2011; Chang et al., 1996; Sohrmann et al., 1996; J.-Q. Wu et al., 2006). Similar precursor structures have been observed in animal cells (Henson et al., 2017; Piekny & Glotzer, 2008). These nodes are positioned in the cell middle by a combination of inhibitory signals from cell ends and positive signals from the cell nucleus (Rincon & Paoletti, 2012, 2016). In the second step, nodes condense to form the CAR, which then undergoes a maturation process. During maturation, many cytokinesis proteins are loaded onto the CAR (Pollard & Wu, 2010; J.-Q. Wu et al., 2003, 2006). In the final step of fission yeast cytokinesis, the CAR constricts in coordination with septum assembly to complete the division process (Pollard & Wu, 2010; Proctor et al., 2012).

The key protein determining position of fission yeast cytokinesis is the anillin-like protein Mid1. Cells without Mid1 assemble off-centered division planes at various angles along the cell surface (Chang et al., 1996; Paoletti & Chang, 2000; Sohrmann et al., 1996; J.-Q. Wu et al., 2003). Mid1 localizes to the nucleus and to cortical nodes prior to CAR assembly (Bähler, Steever, et al., 1998; Paoletti & Chang, 2000; Sohrmann et al., 1996). While at nodes, Mid1 recruits additional cytokinetic ring proteins (Almonacid et al., 2011; Celton-Morizur et al., 2004; Guzman-Vendrell et al., 2013; Motegi et al., 2004; Padmanabhan et al., 2011). Multiple protein kinases phosphorylate Mid1 to regulate its localization. For example, Mid1 is phosphorylated by the polo kinase Plo1 following mitotic entry. This modification triggers the mass release of Mid1 from the nucleus, enabling the protein to accumulate at nodes and recruit downstream proteins (Almonacid et al., 2011; Bähler, Steever, et al., 1998). Subsequently, the NDR family kinase Sid2 phosphorylates Mid1 at the onset of ring constriction and signals for Mid1 to leave the CAR and reaccumulate in the nucleus (Willet et al., 2019a). Other protein kinases that regulate Mid1 include Cdc2/Cdk1, Aurora kinase Ark1, Pdk1, and Pak1/Orb2 (Almonacid et al., 2011; Bimbó et al., 2005; Magliozzi et al., 2020; Rezig et al., 2021, 2023). Despite the wealth of knowledge on kinases that regulate Mid1, far less is known about protein phosphatases. Mid1 binds to the protein phosphatase Clp1, but this interaction positions Clp1 to act on other proteins at the CAR (Clifford et al., 2008). It is also important to note that regulation of Mid1 by protein kinases has been shown to act exclusively on Mid1 localization, whereas other regulatory mechanisms remain unexplored.

In this study, we found that the protein phosphatase PP2A-B56 regulates cytokinesis through Mid1. PP2A functions as a trimeric holoenzyme containing catalytic, scaffolding, and regulatory subunits (Barr et al., 2004). In fission yeast, Par1 is the major isoform of the B56 regulatory subunit (Jiang & Hallberg, 2000). Unlike protein kinases that regulate Mid1 localization, our data show that PP2A-B56 regulates the protein levels of Mid1 for proper cytokinesis. Cells lacking Par1 have reduced levels of Mid1 and exhibit cytokinesis defects including the assembly of off-centered CARs that slide out towards cell tips during maturation. These defects are rescued by restoring Mid1 levels in *par1Δ* cells. Our work defines a new mechanism regulating Mid1 through a conserved protein phosphatase complex.

## RESULTS

### PP2A mutants divide asymmetrically

To identify protein phosphatases that regulate cytokinesis, we screened a set of 9 phosphatase mutant strains for cytokinesis defects. For each strain, cells were stained with Blankophor to mark the division plane and cell outline and then imaged by fluorescence microscopy. Mutants were compared to wild type cells, which had division planes positioned in the cell middle as expected. Most phosphatase mutants maintained this patterning, but *par1Δ*, *ppa2Δ,* and *pzh1Δ* mutants appeared to form off-center division planes (Figure 1A). To quantify this defect, we measured the cell half ratio for each strain examined (Figure 1B). This analysis confirmed that *par1Δ*, *ppa2Δ,* and *pzh1Δ* cells divide asymmetrically compared to wild type (Figure 1C). Par1 is the major B56 regulatory subunit for the PP2A phosphatase in fission yeast (Jiang & Hallberg, 2000), Ppa2 is the major catalytic subunit of PP2A (Jiang & Hallberg, 2001; N. Kinoshita et al., 1993), and Pzh1 is fission yeast protein phosphatase Z (Balcells et al., 1997). Because we identified two mutants in PP2A and the strongest defect was observed for Par1/B56, we investigated this phenotype further.

**Figure 1.**
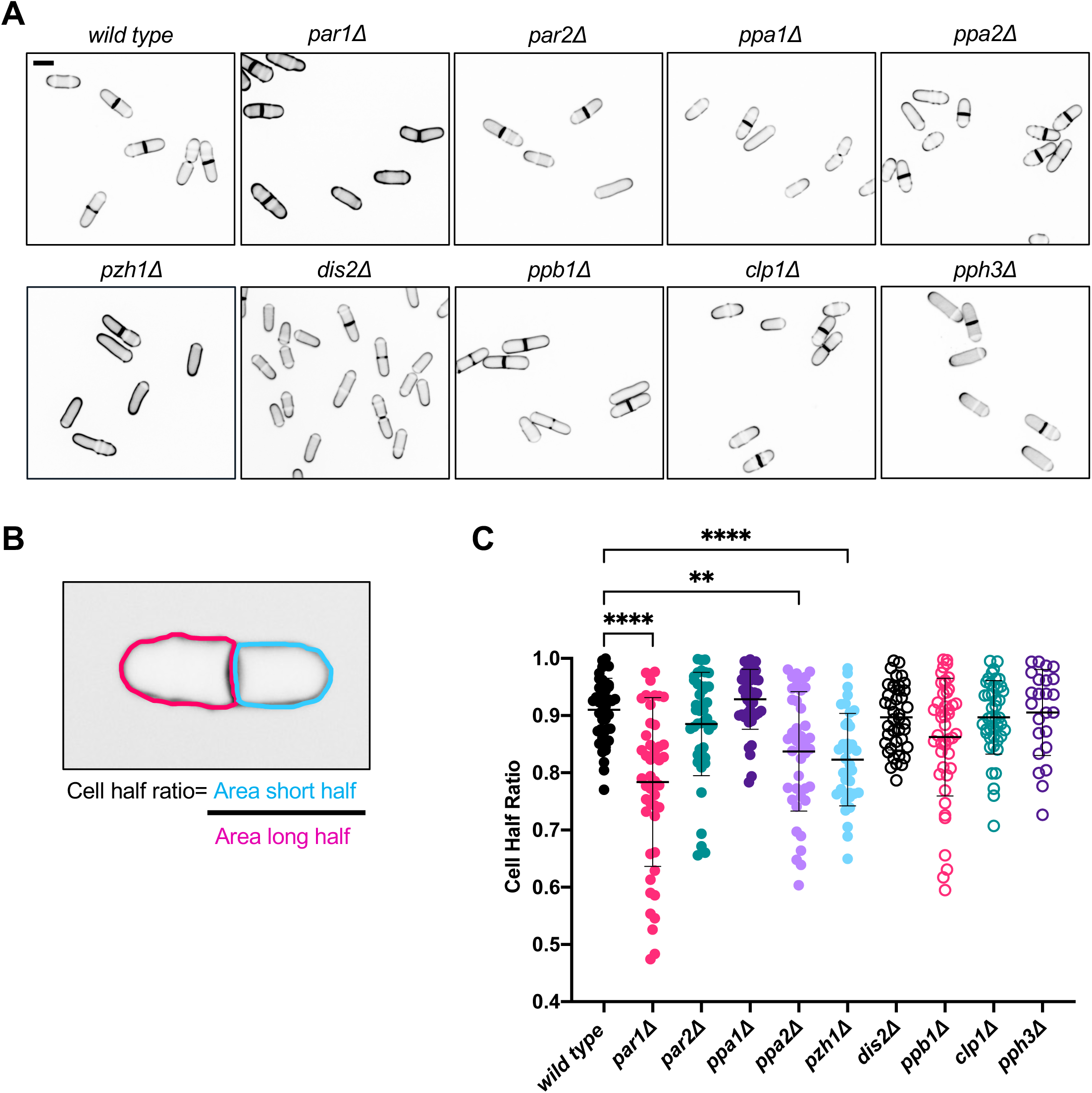
PP2A mutants divide asymmetrically. (A) Representative images of phosphatase mutants stained with Blankophor to mark the cell wall. Images are a single middle focal plane with inverted fluorescence. Scale bar is 7µm. (B) Method for calculating cell half ratio as a measure of division plane symmetry. (C) Cell half ratio for the phosphatase mutants screened. N ≥ 24 cells for each strain. Error bars correspond to the mean ± standard deviation. ** indicates *p* value <0.005; **** indicates *p* value <0.0001; all others are nonsignificant (*p* value >0.05) by one-way analysis of variance (ANOVA) with Dunnett’s multiple comparison test.

The position of the cytokinetic ring determines the position of the division plane, so we monitored the assembly and constriction of cytokinetic rings in *par1Δ* mutants. Using Rlc1-mNeonGreen (mNG) as a marker for the CAR, we found that cytokinesis took longer and resulted in off-center rings for *par1Δ* compared to wild type cells (Figure 2A). Next, we repeated our time-lapse imaging with cells expressing both Rlc1-mNG and Sad1-mEGFP, which marks the spindle pole body (SPB) and provides a clock for the steps of cytokinesis (Nabeshima et al., 1998). We used this approach to quantify defects in both the temporal and spatial control of cytokinesis for *par1Δ* cells. Compared to wild type cells, *par1Δ* cells took slightly longer to assemble the CAR and more than twice as long for CAR maturation, followed by a slower rate of constriction (Figure 2B-D and Supplementary Figure S1A-B). We also measured spatial defects in the site of ring assembly, which was quantified as the cell half ratio at initial CAR assembly (Figure 2E). We tested if this defect resulted from asymmetric position of the nucleus, which provides a spatial cue for CAR (Daga & Chang, 2005; Motegi et al., 2004; Paoletti & Chang, 2000; Tolic-Nørrelykke et al., 2005). However, we found that the nucleus is properly positioned *par1Δ* cells and that the nucleus remains stationary at the cell middle (Supplementary Figure S1C-F). Beyond defects in CAR assembly, we observed CAR rings slide away from the site of assembly in *par1Δ* cells but not in wild type cells (Figure 2A). This sliding of intact CARs occurred during the extended maturation phase and resulted in some CARs constricting over 1 µm away from their assembly site (Figure 2F). We conclude that *par1Δ* cells divide asymmetrically due to the assembly of off-center CARs that slide away from their assembly site. These defects raise the possibility that PP2A-B56 regulates factors involved in the spatial control of CAR assembly and anchoring.

**Figure 2.**
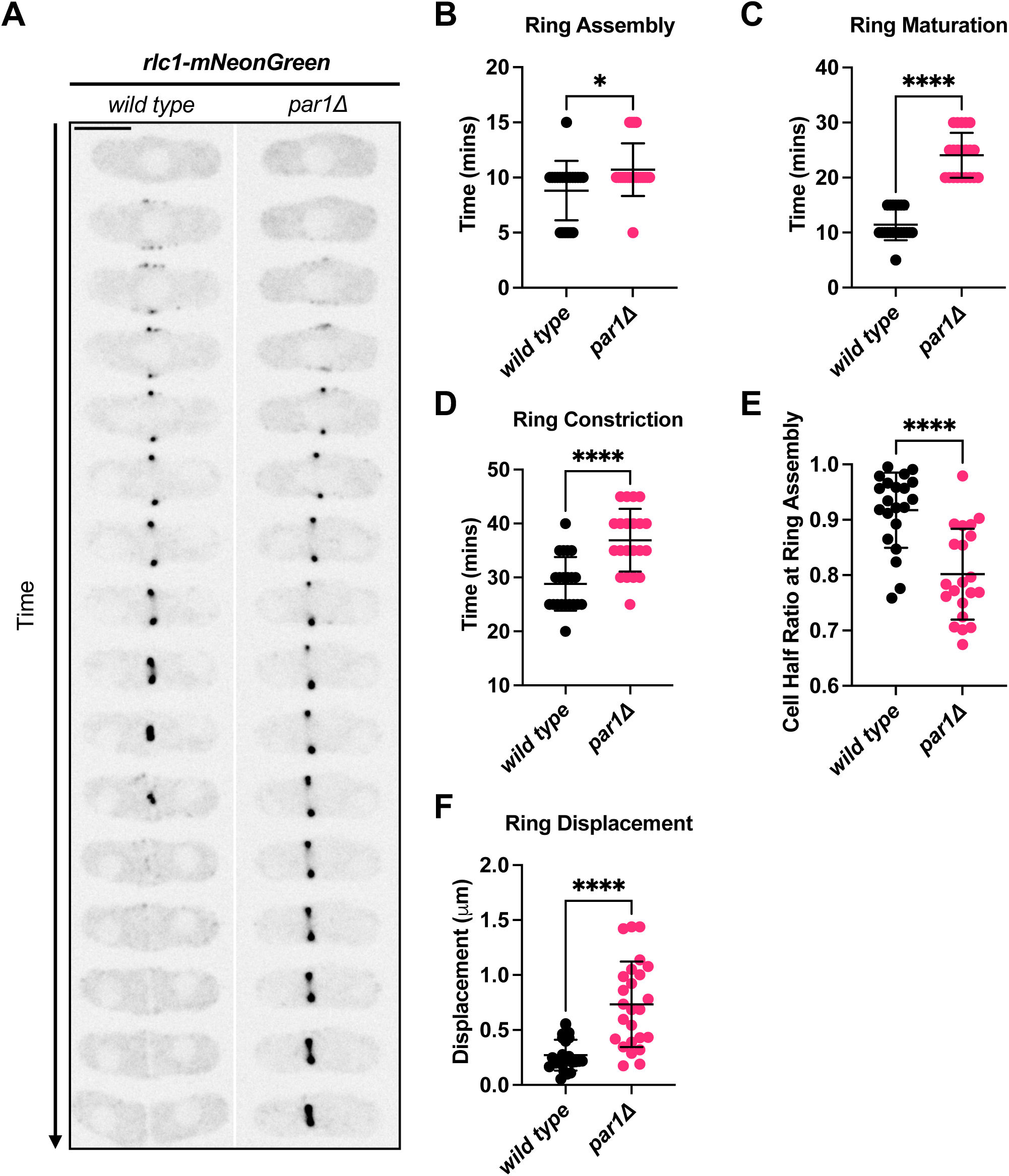
Cytokinetic ring defects in *par1Δ* cells. (A) Representative time-lapse montage of Rlc1-mNeonGreen in wild type and *par1Δ* cells. Images are a single middle focal plane with inverted fluorescence. Images were captured in 5-minute intervals. Scale bar is 5µm. (B-D) Duration of ring assembly (B), ring maturation (C), and ring constriction (D) in wild type and *par1Δ* cells. * Indicates *p* value <0.05; **** indicates *p* value <0.0001. (E) Cell half ratios at time of ring assembly in wild type and *par1Δ* cells. **** indicates *p* value <0.0001. (F) Quantification of cytokinetic ring sliding in wild type and *par1Δ* cells. **** indicates *p* value <0.0001. At least 20 cells were analyzed per strain for all quantifications in this figure. All statistical tests in this figure are Welch’s unpaired t-tests.

### Reduced Mid1 levels cause cytokinesis defects in *par1Δ* cells

To identify the molecular cause of cytokinesis defects in *par1Δ* cells, we tested the localization of a panel of proteins with the potential to anchor CAR position in the cell middle. Several of these factors when mutated lead to ring sliding similar to *par1Δ*. This group includes Cdc15, Rga7, and Bgs1 (Arasada & Pollard, 2014; Martín-García et al., 2014; McDonald et al., 2015; Roberts-Galbraith et al., 2009). We reasoned that reduction of a putative CAR anchor could lead to the observed ring sliding defects in *par1Δ* cells. We tested other factors including Mid1, which acts as a scaffold by binding to both lipids and other CAR proteins, as well as several regulatory protein kinases. For each factor, we measured the fluorescence intensity of its localization at the CAR of wild type versus *par1Δ* cells. Strikingly, Mid1 was the only tested protein with significantly reduced levels at the CAR in *par1Δ* cells (Figure 3A-B and Supplemental Figures S2A-I). All other proteins were either unaffected or increased at the CAR of *par1Δ* cells.

**Figure 3.**
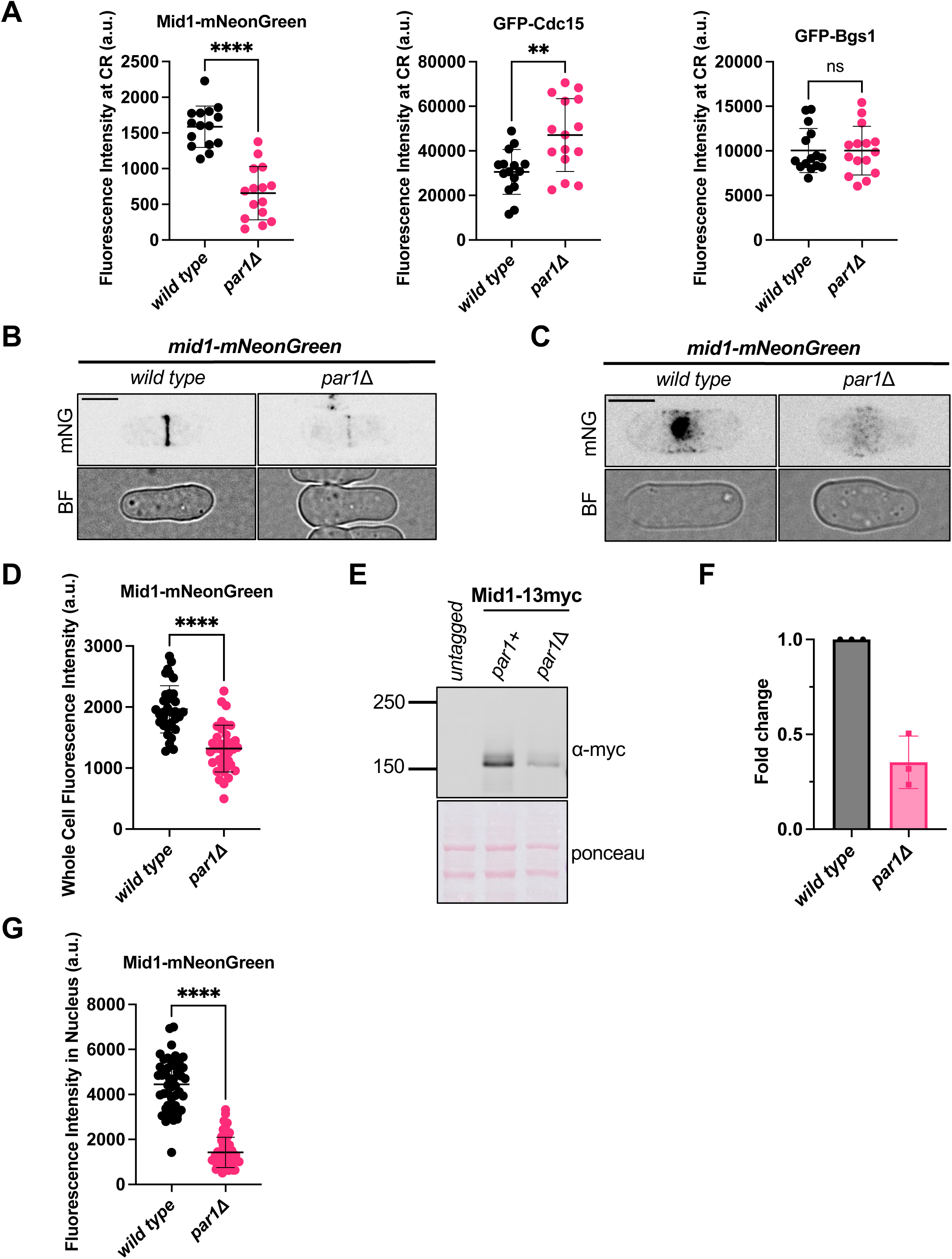
Mid1 levels at CAR are reduced in *par1Δ* cells. (A) Quantification of Mid1-mNeonGreen, GFP-Cdc15, or GFP-Bgs1 at the cytokinetic ring in wild type and *par1Δ* cells. N ≥ 15 cells each. ns indicates *p* value >0.05; ** indicates *p* value <0.005; **** indicates *p* value <0.0001. (B) Representative images of wild type and *par1Δ* cells expressing Mid1-mNeonGreen. Brightfield images are a single middle focal plane. mNG images are sum projections with inverted fluorescence. Scale bar is 5µm. (C) Images of representative wild type and *par1Δ* interphase cells expressing Mid1-mNG. Images are sum projections with inverted fluorescence. Scale bar is 5µm. (D) Quantification of whole cell fluorescence intensity of Mid1-mNG in wild type and *par1Δ* cells. N ≥ 33 cells each. **** indicates *p* value <0.0001. (E) Western blot analysis of Mid1-13myc levels in whole-cell lysates from wild type and *par1Δ* cells. (F) Quantification of western blot analysis of Mid1-13myc levels in wild type and *par1Δ* cells. Ponceau was used to normalize band intensity to total protein. Fold change was determined by taking the ratio of normalized intensity of *par1Δ* to the normalized intensity of wild type. Error bars represent the standard deviation from three independent biological replicates. (G) Quantification of fluorescence intensity of Mid1-mNG in the nucleus in wild type and *par1Δ* cells. N ≥ 48 cells each. **** indicates *p* value <0.0001. All statistical tests in this figure are Welch’s unpaired t-tests.

Reduced Mid1 levels at the CAR of *par1Δ* cells could reflect defective localization of Mid1 protein in these cells, or alternatively a global reduction of Mid1 levels. To distinguish between these possibilities, we measured the whole cell concentration of Mid1 protein in wild type and *par1Δ* cells. By fluorescence microscopy, we observed a significant decrease in Mid1-mNG in *par1Δ* compared to wild type cells (Figure 3C-D and Supplemental Figure S3A). This decrease was confirmed using Western blot of whole cell extracts, which showed reduced Mid1-13myc protein levels in *par1Δ* compared to wild type cells (Figure 3E-F). Previous studies have shown that a major pool of Mid1 resides in the cell nucleus during interphase (Bähler, Steever, et al., 1998; Paoletti & Chang, 2000). We observed reduced nuclear Mid1-mNG signal in *par1Δ* versus wild type cells (Figures 3C and 3G). Together, these results show that *par1Δ* cells have reduced levels of Mid1 protein, resulting in less Mid1 at the CAR. Given the major role of Mid1 in the spatial control of cytokinesis, reduced Mid1 levels might contribute to CAR positioning defects in *par1Δ* cells.

If reduced levels of Mid1 cause the cytokinesis defects in *par1Δ* cells, then restoring Mid1 levels should suppress *par1Δ* cytokinesis defects. We tested this prediction by constructing a *2xmid1-mNG* strain with two copies of Mid1 marked with mNeonGreen. Live-cell imaging confirmed that this approach doubled the Mid1 concentration in both the whole cells (Figure 4A-B and Supplemental Figure S3A) and at the cytokinetic ring (Figure 4C-D). This change in concentration means that *2xmid1-mNG par1Δ* cells have a similar concentration of Mid1 when compared to wild type cells expressing a single copy of *mid1-mNG*. Strikingly, this increased concentration of Mid1 suppressed the cytokinesis defects in *par1Δ* cells. *2xmid1-mNG* significantly improved the cell half ratio of *par1Δ* cells (Figure 4E), and timelapse imaging also revealed a significant reduction of CAR displacement in *2xmid1-mNG par1Δ* cells as compared to *par1Δ* alone (Figure 4F). We note that additional factors are likely to be involved because the suppression is not complete. Nonetheless, this phenotypic suppression indicates that reduced Mid1 levels lead to cytokinesis defects in *par1Δ* cells.

**Figure 4.**
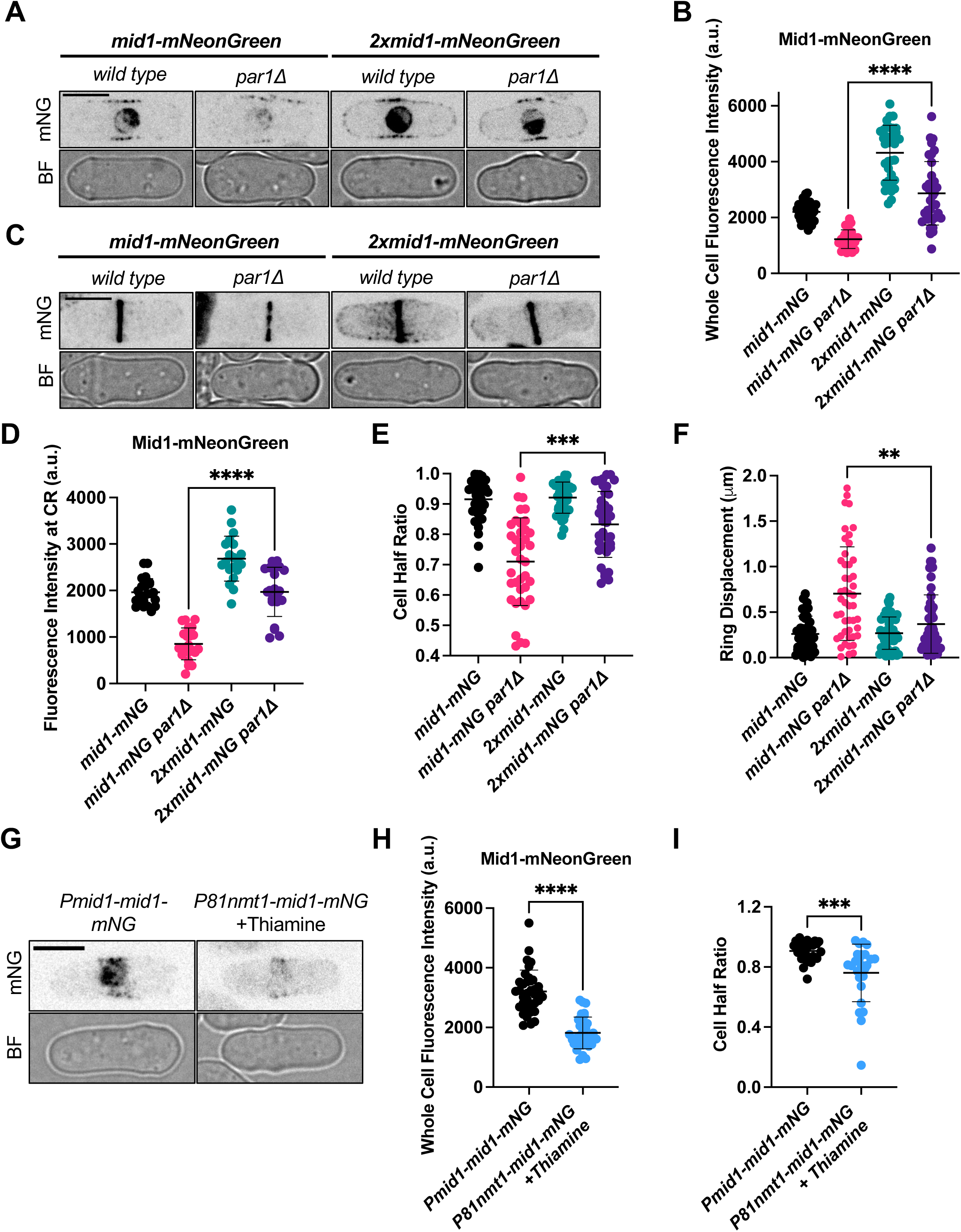
Restoring Mid1 levels suppresses *par1Δ* defects. (A) Representative images of wild type and *par1Δ* strains expressing Mid1-mNeonGreen (left) or 2xMid1-mNeonGreen (right). Brightfield images are a single middle focal plane. mNG images are a single middle focal plane with inverted fluorescence. Scale bar is 5µm. (B) Quantification of whole cell fluorescence intensity of Mid1-mNG in the indicated strains. Whole cell fluorescence intensity was determined from sum intensity projections. N ≥ 35 cells each. (C) Representative images of wild type and *par1Δ* cells expressing Mid1-mNG (left) or 2xMid1-mNG (right). Brightfield images are a single middle focal plane. mNG images are sum projections with inverted fluorescence. Scale bar is 5µm. (D) Quantification of the fluorescence intensity of Mid1-mNG at the cytokinetic ring in the indicated strains. N ≥ 20 cells each. (E) Cell half ratio for the indicated strains. N ≥ 35 cells each. (F) Quantification of cytokinetic ring displacement in the indicated strains. N ≥ 46 cells each. (G) Representative images the indicated strains. *P81nmt1-mid1-mNG* cells were grown in 0.005μg/ml thiamine. Brightfield images are a single middle focal plane. mNG images are sum projections with inverted fluorescence. Scale bar is 5µm. (H) Quantification of whole cell fluorescence intensity of Mid1-mNG in the indicated strains. Whole cell fluorescence intensity was determined from sum projections. N ≥ 32 cells each. (I) Cell half ratio for the indicated strains. N ≥ 25 cells each. Error bars correspond to the mean ± standard deviation. All statistical tests performed in this figure are one-way analysis of variance (ANOVA) with Dunnett’s multiple comparison test. ** Indicates *p* value <0.005; *** indicates *p* value <0.0005; **** indicates *p* value <0.0001; all others are non-significant (*p* value >0.05).

Next, we tested if reducing Mid1 levels in otherwise wild type cells recapitulates the asymmetric cell division observed in *par1Δ* cells. We constructed a strain in which the endogenous *mid1+* promoter was replaced by the thiamine-repressible *P81nmt1* promoter (Basi et al., 1993; Maundrell, 1990). In the absence of thiamine, these *P81nmt1-mid1-mNG* cells exhibited strong expression of Mid1-mNG and medial cell division (Supplemental Figure S3B). Addition of high thiamine (2.5μg/ml) repressed Mid1-mNG expression, resulting in *mid1Δ* phenotypes (Supplemental Figure S3B). By testing a range of thiamine concentrations, we found that addition of 0.005 µg/ml thiamine caused ∼50% reduction in Mid1-mNG protein levels (Figure 4G-H and Supplemental Figure S3C), similar to the effects observed in *par1Δ* cells. This reduction in Mid1-mNG protein levels caused asymmetric division as measured by the cell half ratio (Figure 4I). This result shows that reduced Mid1 levels alone are sufficient to cause asymmetric cell division.

Based on our identification of Mid1 regulation by PP2A combined with past work showing extensive regulation of Mid1 by phosphorylation, we tested if other protein phosphatase mutants also alter Mid1 levels. We quantified the whole cell fluorescence intensity of Mid1-mNG in the same phosphatase mutants that were tested for cytokinesis defects in Figure 1. The only mutants with significantly reduced Mid1 levels were *par1Δ* and *ppa2Δ*, both components of PP2A-B56. All other mutants did not impact Mid1 levels or modestly increased Mid1 levels (Figure 5A-B and Supplemental Figure S4). We conclude that regulation of Mid1 protein levels is specific to PP2A-B56 and does not reflect a general phenotype of protein phosphatase mutants.

**Figure 5.**
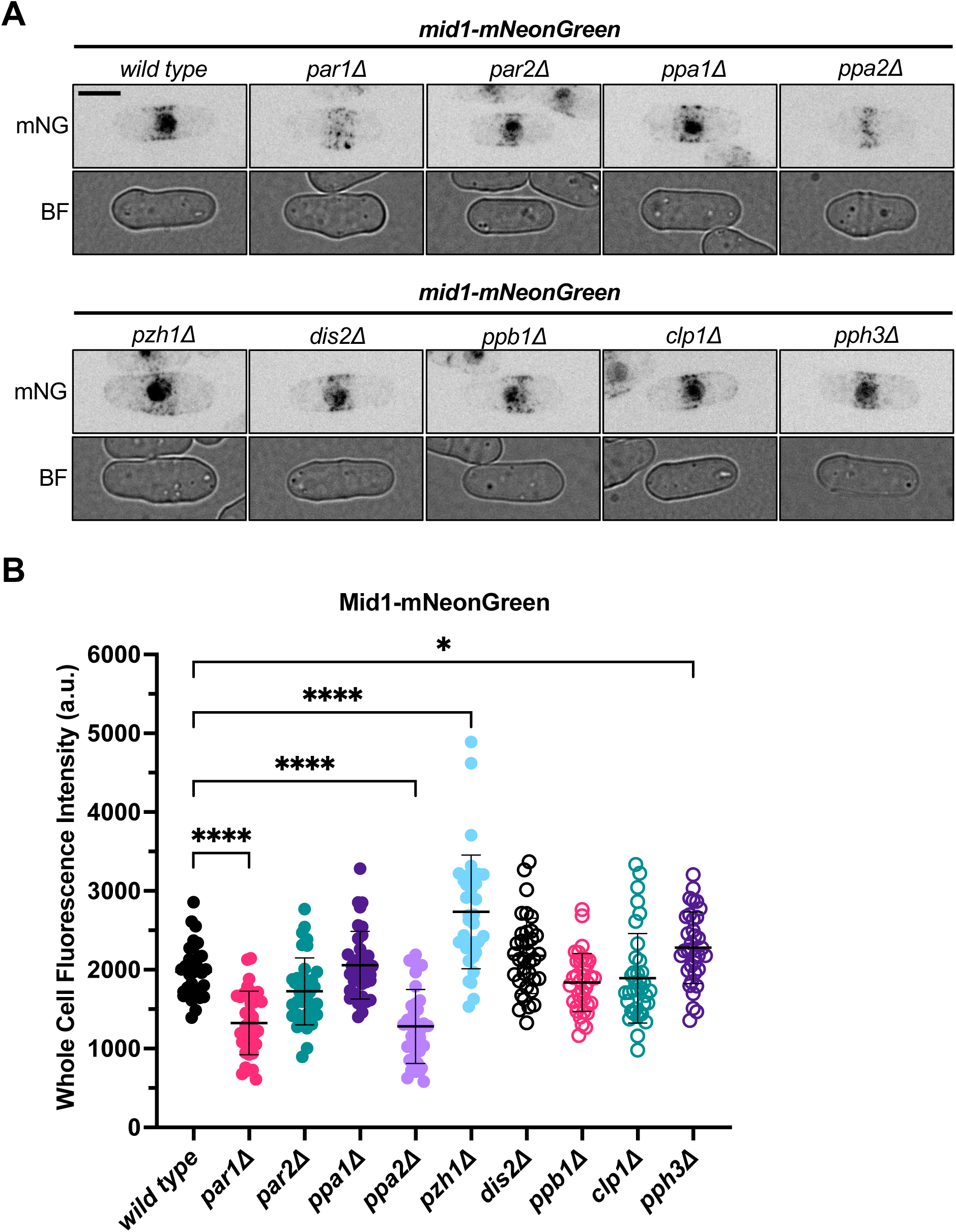
Reduced Mid1 levels are not found in other phosphatase mutants. (A) Representative images of phosphatase mutants expressing Mid1-mNG. Brightfield images are a single middle focal plane. mNG images are sum projections with inverted fluorescence. Scale bar is 5µm. (B) Quantification of whole cell fluorescence intensity of Mid1-mNG for the indicated strains. N ≥ 33 cells each. * Indicates *p* value <0.05; **** indicates *p* value <0.0001 by one-way ANOVA with Dunnett’s multiple comparison test. All others are non-significant (*p* value >0.05).

### Par1 localizes to cortical nodes

Mid1 localizes to cytokinetic nodes and to the CAR prior to its constriction. Previous studies have shown Par1 localization to the SPB and the CAR during division (Jiang & Hallberg, 2000; Le Goff et al., 2001). We decided to reexamine Par1 localization based on its regulation of Mid1. We combined Par1-mNG with markers for the CAR (Rlc1-mRuby) and SPB (Sad1-mCherry) in the same strain. Our imaging confirmed Par1 colocalization with SPB and CAR (Figure 6A). We found that Par1 associated with the CAR during maturation and constriction phases (Figure 6B-C). In addition, we observed faint Par1 localization to cytokinetic nodes prior to CAR formation. These cortical Par1 puncta colocalized with Rlc1 nodes (Figure 6D-E), which are known to contain Mid1. We observed Par1 node localization in 9/12 cells that contained Rlc1 nodes but not earlier in the cell cycle. To test if Mid1 regulates Par1 localization to these structures, we imaged Par1-mNG and Rlc1-mCherry in *mid1Δ* cells. In the absence of Mid1, nodes were absent but Par1 still localized to the SPB and CAR (Figure 6F). We conclude that Par1 and Mid1 overlap at nodes and the assembled CAR, which could be sites for Par1/PP2A to regulate Mid1 protein levels.

**Figure 6.**
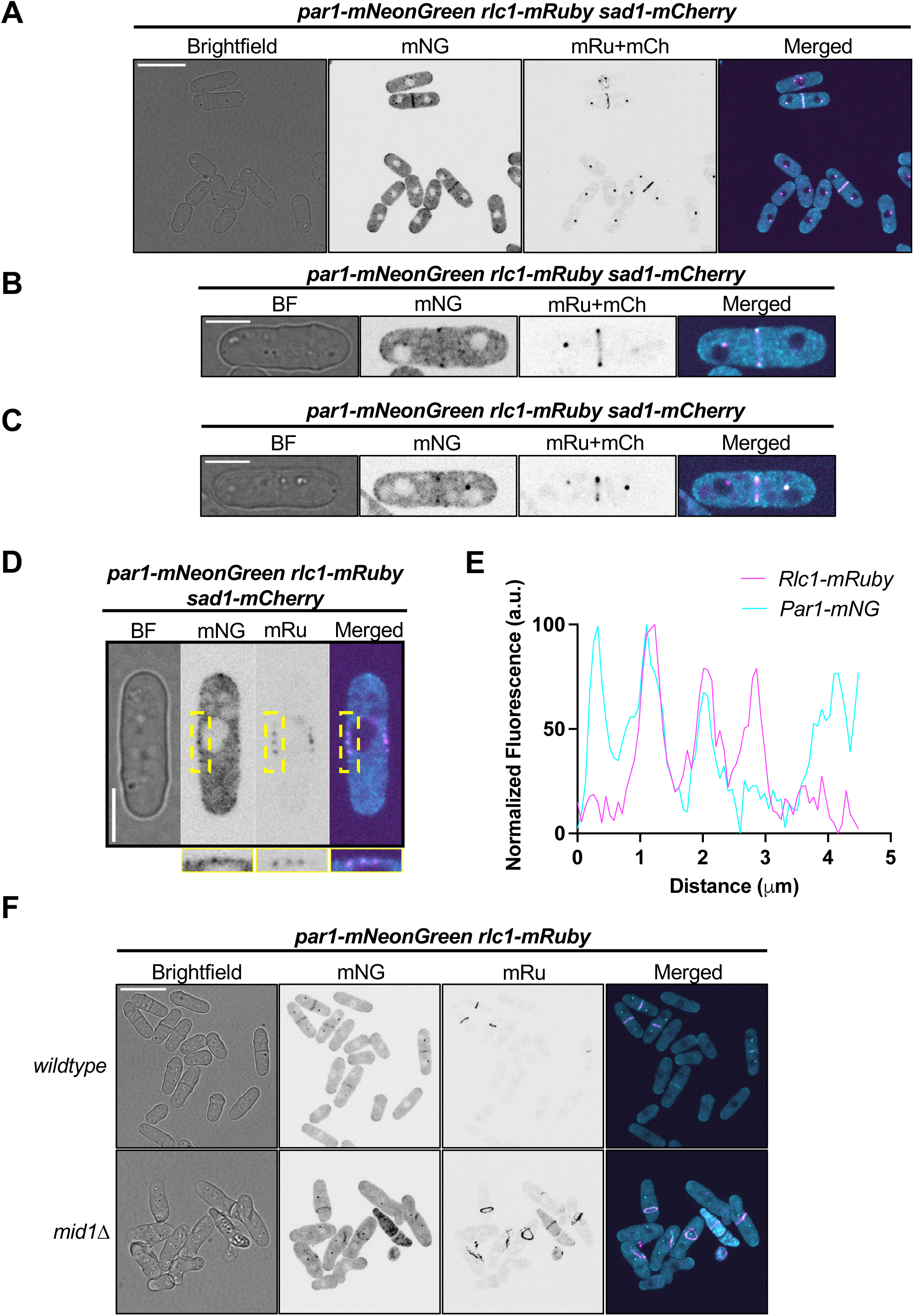
Par1 localizes to nodes and the CAR. (A) Representative images of cells expressing Par1-mNeonGreen, Rlc1-mRuby, and Sad1-mCherry. Brightfield image is a single middle focal plane. mNG and mRuby+mCherry images are maximum intensity projections with inverted fluorescence. Merged image is a maximum intensity projection of mNG and mRu+mCh channels overlaid. Scale bar is 14 µm. (B) Representative cell with a fully assembled CAR. Single middle focal plane images are shown. mNG and mRu+mCh images are with inverted fluorescence. Scale bar is 5 µm. (C) Representative cell with partially constricted CAR. Single middle focal plane images are shown. mNG and mRu+mCh images are with inverted fluorescence. Scale bar is 5 µm. (D) Single middle focal plane images of cells expressing Par1-mNG, Rlc1-mRu, and Sad1-mCh. Scale bar is 5µm. Boxes below show enlarged images of region outlined with a yellow dashed box. (E) Line scans of the boxed region from the merged composite of Par1-mNG (cyan) and Rlc1-mRu (magenta). (F) Representative images of wild type and *mid1Δ* cells expressing Par1-mNG and Rlc1-mRu. Brightfield images are a single middle focal plane. mNG and mRu images are maximum intensity projections with inverted fluorescence. Merged images are maximum intensity projections of mNG and mRu channels overlaid. Scale bar is 14 µm.

### Response of *par1Δ* cells to nuclear displacement

Mid1 shuttles between the nucleus and the adjacent plasma membrane in interphase cells (Almonacid et al., 2009; Bähler, Steever, et al., 1998; Daga & Chang, 2005). The nuclear shuttling of Mid1 allows the nucleus to specify the position of the cytokinetic ring (Almonacid et al., 2009). Because we detected significantly reduced nuclear pools of Mid1, we next asked whether the communication between the nucleus and the ring was impaired in *par1Δ* cells. To test this possibility, we displaced the nucleus by treating cells with the microtubule depolymerizing drug methyl benzimidazole-2-yl-carbamate (MBC) followed by centrifugation (Figure 7A) (Almonacid et al., 2009; Daga & Chang, 2005; Opalko et al., 2023). These cells expressed Cut11-mCherry to mark the nucleus and Rlc1-mNG to label the CAR. After centrifugation, we performed time-lapse imaging of cells maintained in MBC to prevent nuclear movement. CARs formed and constricted over the displaced nuclei in wild type cells, resulting in ‘cut’ nuclei (Figure 7B). In contrast, many *par1Δ* cells with displaced nuclei exhibited CARs that assembled and constricted in the cell middle, despite the absence of the nucleus at this site (Figure 7B). Quantification of nuclear and CAR positions demonstrated that *par1Δ* cells exhibit this defect when the nucleus is partially displaced from the cell middle but not after extreme movement to the cell end (Figure 7C). It is important to note that we measured CAR position based on the initial assembly site to avoid complications from CAR sliding in *par1Δ* mutants.

**Figure 7.**
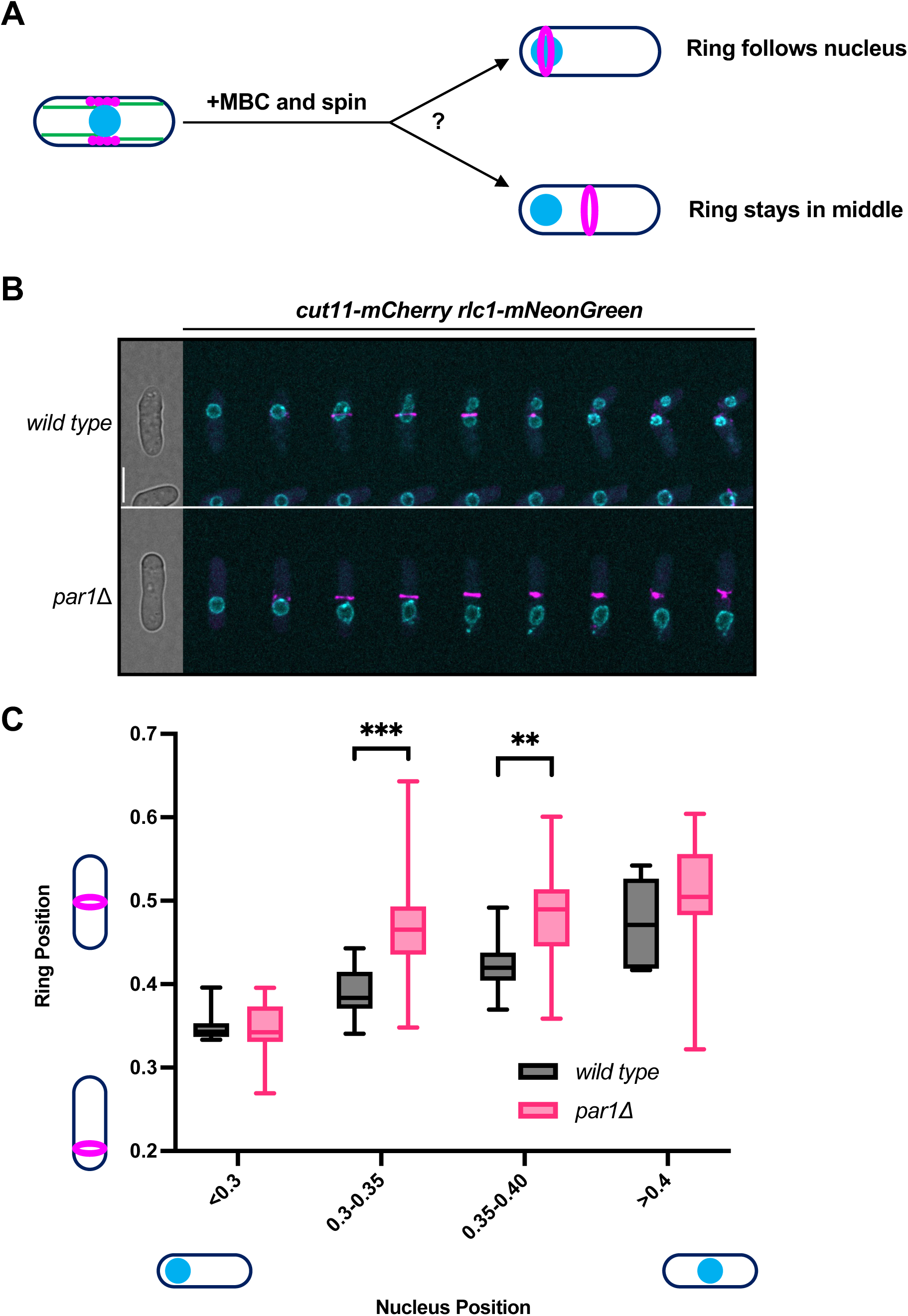
*par1Δ* mutants exhibit partial defects in repositioning the division plane after nuclear displacement. (A) Cartoon outlining nuclear displacement experiments. Cells were treated with 25µg/ml MBC before centrifugation to displace the nucleus. Cells were maintained in MBC, and the positions of the cytokinetic ring and displaced nucleus were examined by fluorescence microscopy. (B) Time-lapse montage of wild type and *par1Δ* cells expressing Cut11-mCherry and Rlc1-mNeonGreen following MBC treatment and centrifugation. Images were captured in 15-minute intervals. Representative maximum intensity projections are shown with Cut11-mCh in cyan and Rlc1-mNG in magenta. Brightfield image is a single middle focal plane from the first time point shown. Scale bar is 7µm. (C) Quantification of nuclear displacement experiments. Positions are expressed as a fraction of the cell length. Nuclear positions were separated into <0.3, 0.3-0.35, 0.35-0.40, and >0.4 bins. Error bars correspond to minimum and maximum ring position for each bin. N ≥ 56 cells each.

If the reduced levels of Mid1 contribute to this defect, then restoring the levels of Mid1 in a *par1Δ* mutant should rescue the ability of the nucleus to specify the position of the division plane. We tested this possibility using *par1Δ 2xmid1-mNG* cells. In this strain, CARs formed and constricted over displaced nuclei similar to wild type cells (Supplemental Figure S5). These results show that *par1Δ* cells have defects in the ability of the nucleus to specify the position of the division plane due to reduced levels of Mid1 protein.

### B56 SLiMs in Mid1

The B56 regulatory subunit of PP2A binds to short linear motifs (SLiMs) in substrates, or in proteins that are close proximity to substrates (Hertz et al., 2016; Kruse et al., 2020; J. Wang et al., 2016; X. Wang et al., 2016; C.-G. Wu et al., 2017). Using the SLiMSearch4 tool, we identified three motifs in Mid1 that fit the B56 SLiM sequence consensus [LMFI]xx[IVL]xE (Figure 8A). We generated a *mid1-3xSLiM* mutant by mutating the three core amino acid residues within each motif to alanine, which is predicted to abolish interactions with Par1/B56. This Mid1-3xSLiM-mNG mutant displayed a small but significant reduction in expression level compared to wild type Mid1-mNG (Figure 8B-C). The reduction was not as dramatic as *par1*Δ and did not lead to significant defects in division plane positioning (Figure 8D-E) or the ability of displaced nuclei to specify the division plane (Supplemental Figure S6B).

**Figure 8.**
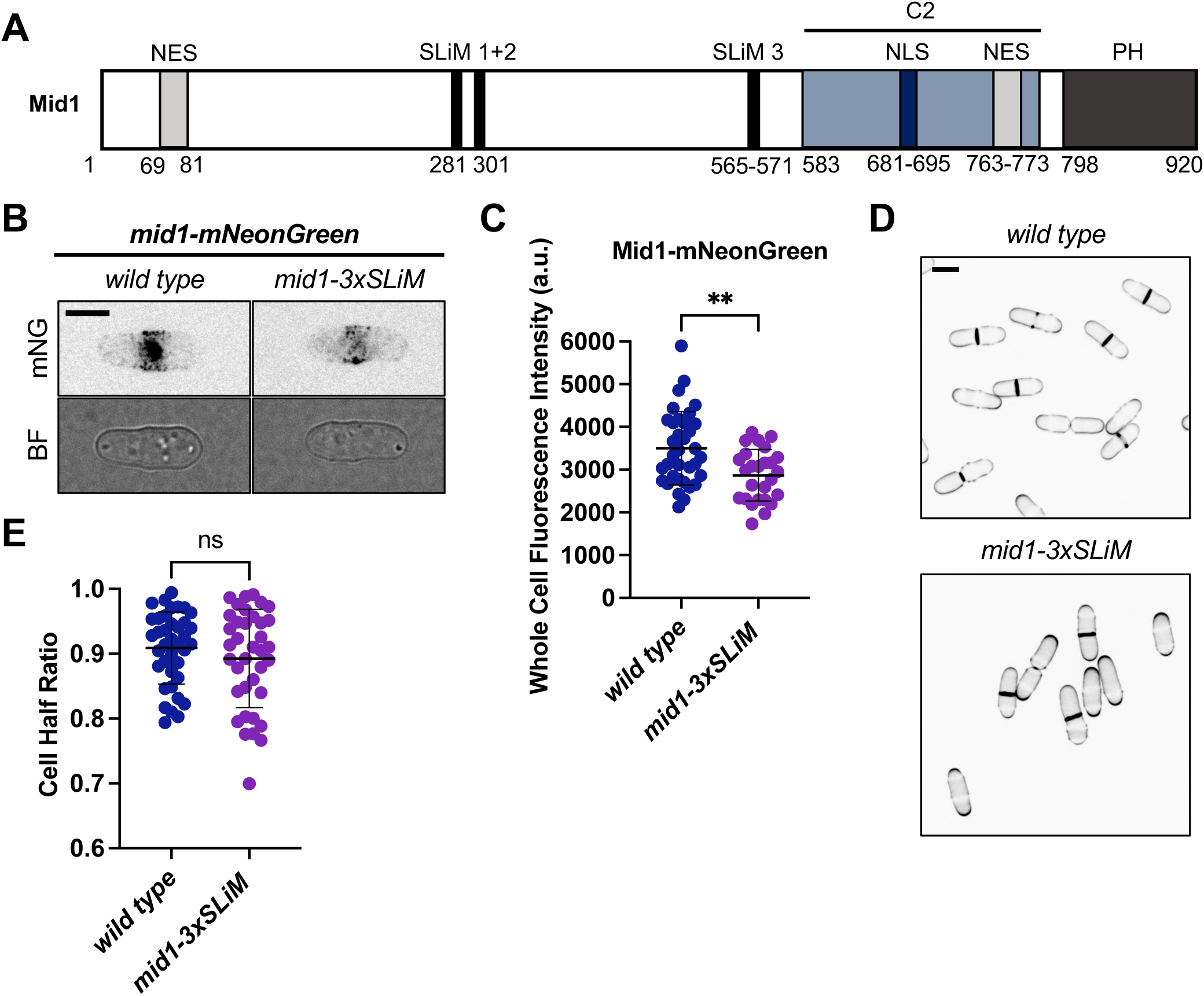
Mutation of Mid1 SLiMs does not lead to asymmetric divisions. (A) Mid1 domain organization including three LxxIxE short linear motifs that represent potential PP2A-B56 interaction sites. NES, nuclear export sequence. SLiM, short linear interaction motif. NLS, nuclear localization sequence. PH, pleckstrin homology. (B) Representative wild type and *mid1-3xSLIM* cells expressing Mid1-mNG. Brightfield images are a single middle focal plane. mNG images are sum projections with inverted fluorescence. Scale bar is 5µm. (C) Quantification of whole cell fluorescence intensity of Mid1-mNG in wild type and *mid1-3xSLIM* cells. N ≥ 25 cells each. ** indicates *p* value <0.005. (D) Representative images of wild type and *mid1-3xSLIM* cells stained with Blankophor to mark the cell wall. Images are a single middle focal plane with inverted fluorescence. Scale bar is 5µm. (E) Quantification of cell half ratios for wild type and *mid1-3xSLIM*. N ≥ 36 cells each. ns indicates *p* value >0.05.

To further investigate the effects of *mid1-3xSLiM*, we tested for synthetic genetic interactions with other cytokinesis mutants including *pom1Δ*, which causes misplaced CARs and septa (Bähler & Pringle, 1998), and *rng2-D5*, a temperature-sensitive mutant that impairs the formation of CARs (Chang et al., 1996; Eng et al., 1998). We observed synthetic growth defects for both *mid1-3xSLiM pom1Δ* and *mid1-3xSLiM rng2-D5* double mutants (Figure 9A). In addition, these double mutants exhibited division plane defects that were more severe than the single mutants. These defects included an increased frequency in tilted septa (Figure 9B-C). Taken together, these genetic results identify a role for B56 SLiM motifs in Mid1 in promoting proper cytokinesis. We note that the 3xSLiM mutation does not impair Mid1 protein levels or cytokinesis to the same degree as *par1Δ*. One potential explanation is that many additional proteins at cortical nodes and the cytokinetic ring also contain the same [LMFI]xx[IVL]xE motif (Supplementary Table S2), meaning that PP2A-B56 could potentially target Mid1 by interacting with these proximal proteins in addition to any role for the Mid1 SLiMs.

**Figure 9.**
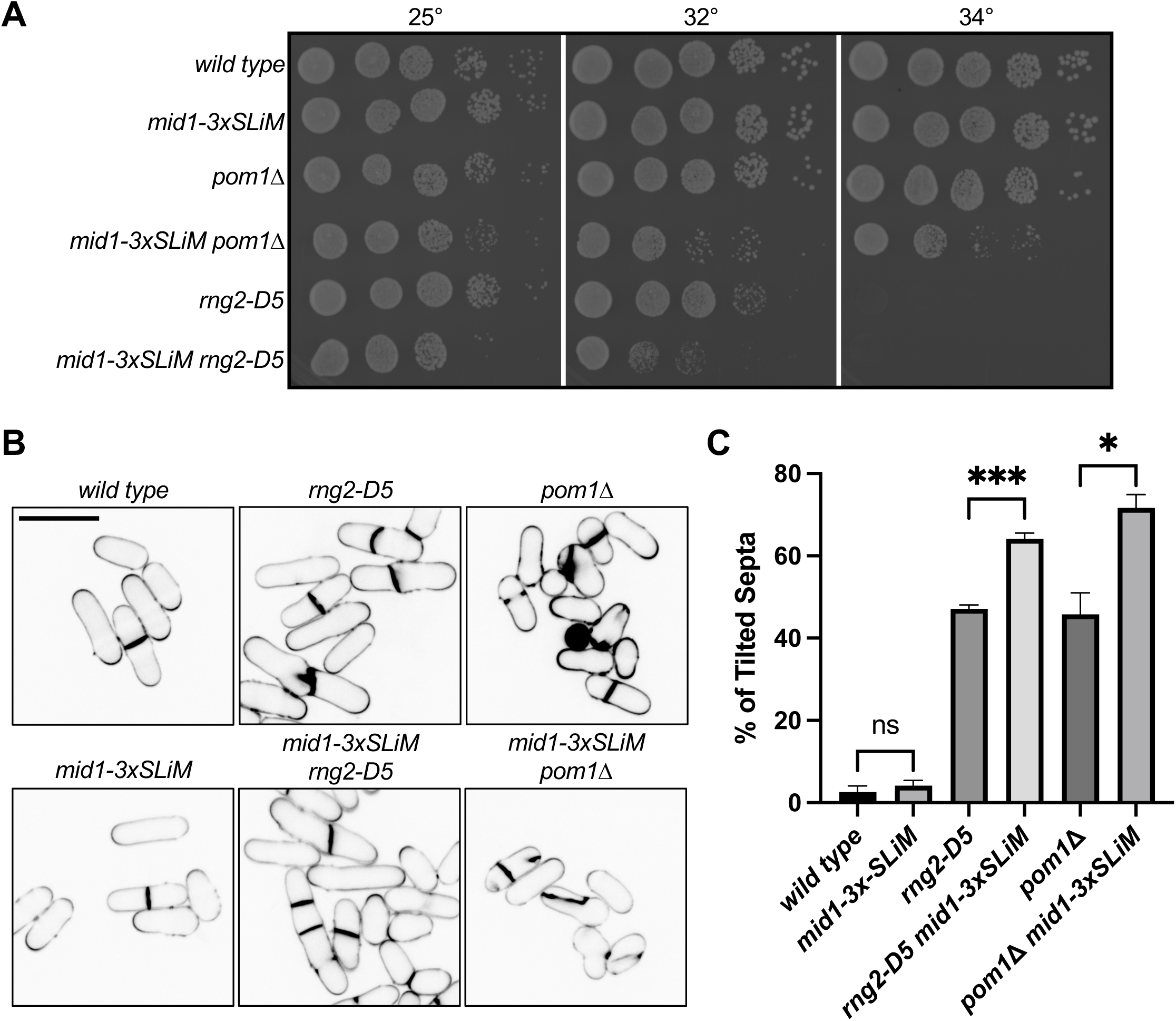
Synthetic genetic defects between *mid1-3xSLiM* mutant and cytokinesis mutants. (A) Serial-dilution growth assays for each indicated strain at 25° (left), 32° (middle), and 34° (right). (B) Representative images of cells stained with Blankophor to mark the cell wall for the indicated strains. Images are a single middle focal plane with inverted fluorescence. Scale bar is 14µm. (C) Quantification of division plane defects for the indicated strains. Values are means ± standard deviation from three biological experiments. N ≥ 55 cells each.

## DISCUSSION

In this study, we found that the protein phosphatase PP2A promotes efficient cytokinesis through Mid1. In the absence of Par1, the major B56 regulatory subunit of PP2A in fission yeast, cells have reduced levels of cellular Mid1 leading to cytokinetic defects. These defects include contractile rings that form off-centered and slide out towards cell tips during ring maturation. Restoring the levels of Mid1 in *par1Δ* cells can rescue the cytokinetic defects, thereby establishing a connection between PP2A-B56, Mid1 abundance, and cytokinesis efficiency. Mid1 is known to be a highly phosphorylated protein and previous groups have shown that phosphorylation dictates Mid1 subcellular localization throughout the cell cycle (Almonacid et al., 2011; Bähler, Steever, et al., 1998; Willet et al., 2019b). Our work has identified a new mode of regulation by which protein phosphorylation regulates Mid1 abundance to promote cytokinesis.

Reversible protein phosphorylation regulates many aspects of cell division including cytokinesis (Barr et al., 2011; Nilsson, 2019; Trinkle-Mulcahy & Lamond, 2006). While understudied relative to their kinase counterparts, protein phosphatases are increasingly appreciated as key regulators of cytokinesis (Bollen et al., 2009). For instance, the loss of PP2A and its regulators causes cytokinetic defects in both budding and fission yeast (Goyal & Simanis, 2012; Healy et al., 1991; Jiang & Hallberg, 2001; K. Kinoshita et al., 1996; Le Goff et al., 2001; Moyano-Rodríguez et al., 2022; Moyano-Rodriguez & Queralt, 2019; Schutt & Moseley, 2020). Budding yeast cells lacking Rts1, the B56 regulatory subunit of PP2A, are multinucleated and elongated at elevated temperatures, suggesting a failure to complete cytokinesis (Moyano-Rodriguez & Queralt, 2019). Similarly, deletion of both fission yeast B56 subunits Par1 and Par2 lead to multiple cytokinesis defects (Jiang & Hallberg, 2000). Results from our study provide an explanation for these defects through the cytokinesis regulator Mid1. We note that *par1Δ* mutants exhibit defects in cell size and cell polarity (Jiang & Hallberg, 2001), which can influence nuclear positioning and cytokinesis (Ashraf et al., 2021; Mishra et al., 2012), so additional factors likely work with Mid1 to link Par1 with cytokinesis.

We only observed reduced Mid1 protein levels in cells lacking Par1 but not Par2. This result can be explained by previous estimations that Par1 accounts for more than 90% of the B56 regulatory subunits in *S. pombe* (Jiang & Hallberg, 2000). We also note that Par1 and Par2 have distinct subcellular localizations (Jiang & Hallberg, 2000; Le Goff et al., 2001). These findings connect with our previous observation that mutants in fission yeast Sds23, a poorly understood regulator of PP2A and PP6 phosphatases, have defects in Mid1 levels and cytokinesis (Schutt & Moseley, 2020). These results follow a general theme that tightly controlled formation and regulation of specific PP2A holoenzymes is required to control cell division events in time and space (Peris et al., 2023).

PP2A is one of the most abundant enzymes and its activity constitutes the majority of Serine/Threonine phosphatase activity in any given cell (Janssens & Goris, 2001). In humans, the B56 regulatory family is encoded by five genes (Seshacharyulu et al., 2013). These isoforms demonstrate distinct subcellular localization and appear to regulate distinct processes (Vallardi et al., 2019). The B56γ and B56ε isoforms seem to be important for cytokinesis because their depletion leads to binucleated HeLa cells (Bastos et al., 2014). Consistent with this finding, cytokinesis failed to progress normally when B56β/ B56γ/ B56ε were depleted in HeLa cells (C.-G. Wu et al., 2017). These cytokinetic defects have been linked to the development of aneuploidy, which is a common characteristic of cancer (Weaver et al., 2007). These findings indicate that PP2A-B56 likely plays a conserved role in cytokinesis and our results in fission yeast could point to new mechanistic roles in other cells.

The B56 regulatory subunit determines substrate specificity of PP2A by recognizing short linear motifs (SLiMs) in disordered regions of interacting proteins (Hertz et al., 2016; Kruse et al., 2020; J. Wang et al., 2016; X. Wang et al., 2016; C.-G. Wu et al., 2017). Interestingly, our *mid1-3xSLIM* mutant failed to phenocopy the cytokinetic defects that we observed in *par1Δ* cells, and we have not detected a physical interaction between Par1 and Mid1. Therefore, we envision two potential models for PP2A-Par1 regulation of Mid1 and cytokinesis. First, PP2A-Par1 might bind to another protein that allows it to act on Mid1. This possibility is supported by examples of proteins such as BubR1 that contain SLiMs and serve as “docking” factors for PP2A-B56 to act on other substrates (Suijkerbuijk et al., 2012), as well as other cortical node proteins with predicted SLiMs for B56 interaction. A second model is that PP2A does not act directly on Mid1 but rather regulates proteins that control Mid1 protein levels through synthesis and/or degradation. These two models are not mutually exclusive and will be addressed in future studies. We also note a potential role for PP2A-B56 in regulating the nuclear shuttling of Mid1 based on its absence from nuclei in *par1Δ* cells. Beyond Mid1, *par1Δ* also altered the levels of other cytokinesis proteins such as Cdc15 at the CAR. It will be interesting to learn if PP2A-Par1 uses similar or distinct mechanisms to regulate these different proteins.

We found that cytokinesis defects arise from decreased Mid1 levels, and previous work has shown that increased Mid1 levels also cause cytokinesis defects (Paoletti & Chang, 2000). This requirement for proper Mid1 expression could represent a conserved principle because altered levels of the Mid1-related protein anillin in higher eukaryotes similarly result in cytokinetic failure. For example, depletion of anillin by RNA interference (RNAi) leads to cytokinetic failure in *Drosophila* (Echard et al., 2004; Hickson & O’Farrell, 2008; Somma et al., 2002; Straight et al., 2005). The same failure to complete cytokinesis was observed in human cells following the knockdown of anillin (Straight et al., 2005; Zhao & Fang, 2005). Further, dysregulation of anillin levels contributes to the tumorigenesis of numerous hematologic and solid tumors (Uhlén et al., 2015), and anillin overexpression has been linked to poor patient outcomes in breast cancer (Dai et al., 2019; Hall et al., 2005; Magnusson et al., 2016; D. Wang et al., 2020; Z. Wang et al., 2017; Zhou et al., 2015). The knockdown of anillin in breast cancer cells attenuates migration and cancer progression, further solidifying the role of the protein in tumorigenesis (Magnusson et al., 2016; Zhou et al., 2015). Future work on the pathways that regulate expression levels of Mid1 and anillin will provide insight into mechanisms that ensure proper cytokinesis.

## METHODS

### Strain construction and growth

Standard *Schizosaccharomyces pombe* media and methods were used (Moreno et al., 1991). Strains used in this study are listed in Supplementary Table S1. Gene deletions and tagging were performed using PCR and homologous recombination (Bähler, Wu, et al., 1998).

*Pmid1-mid1-mNeonGreen-Tadh1* was generated by PCR from genomic DNA of JM5288 and inserted into pJK210, as previously described (Magliozzi et al., 2020). The resulting *pJK210-Pmid1-mid1-mNeonGreen-Tadh1* plasmid was linearized by digestion with StuI and integrated into the genomic *ura4-294* locus of JM1643 (*ura4-294 leu1-32 ade6-M210 h+)*. The resulting strain was crossed to JM6994 (*mid1-mNeonGreen::hphR h-)* to generate JM7912 (*ura4+::[pJK210-PMid1-mid1-mNeonGreen-Tadh1] mid1-mNeonGreen::hphR leu? ade? h-*) used in Figure 4. Hygromycin resistant progeny were screened by microscopy and the whole cell fluorescence intensity of Mid1-mNG was determined to confirm 2xMid1-mNG.

*Pmid1-mid1-mNeonGreen-Tadh1* was ligated into pDC99 using KpnI and SacII sites. The *mid1-3xSLiM* mutant was synthesized as a GBlock (Integrated DNA Technologies) and inserted into the pDC99 plasmid by Gibson assembly (New England Biolabs). For the *mid1-3xSLiM* mutant, the following residues were mutated to alanine: L281, I284, E286, F296, V299, E301, L565, V568, and E570. The pDC99 plasmids used in this study were linearized by PCR and integrated into the *leu1* locus of JM737 (*ura4-D18 h+).* Correct colonies were those that grew on EMM-Uri medium but not on EMM-Leu medium. The resulting strains were crossed to JM8072 (*mid1Δ::kanMX6 leu+ ura4-D18 h-*) to eliminate the copy at the endogenous *mid1* locus. All plasmids were verified by Sanger sequencing.

*pDC99-P81nmt1-mid1-mNeonGreen-Tadh1* was generated by Gibson assembly using PCR products. The *P81nmt1* insert was amplified off of pJM383 (*pUC19-nmt81::ura4+*). The backbone fragment was amplified from *pDC99-Pmid1-mid1-mNeonGreen-Tadh1*. The resulting strain was linearized by PCR and integrated into the *leu1* locus of JM737 (*ura4-D18 h+*). Correct colonies were those that grew on EMM-Uri medium but not EMM-Leu medium. The resulting strains were crossed to JM8072 (*mid1Δ::kanMX6 leu+ ura4-D18 h-*) to generate strains where the only copy of mid1 was expressed by *P81nmt1* promoter.

In Figure 9, single and double mutants were constructed by crossing either JM8117 (*mid1Δ::kanMX6 pDC99:Pmid1-mid1-mNeonGreen-Tadh1*) or JM8125 (*mid1Δ::kanMX6 pDC99:Pmid1-mid1(L281A, I284A, E286A, F296A, V299A, E301A, L565A, V568A,*

*E570A)-mNeonGreen-Tadh1*) to known cytokinetic mutants including, *pom111*, *rng2-D5*, *cdc4-31*, *cdc15-140*, or *plo1-1*. We did not observe synthetic genetic interactions between the *mid1-3xSLiM* mutant and *cdc4-31*, *cdc15-140*, or *plo1-1*. In Figure 9A, cells were spotted by 10-fold serial dilutions on YE4S plates and incubated at 25°C, 32°C, or 34°C for three days before imaging. Cells were grown in YE4S at 25°C before being stained with Blankophor to mark the cell wall in Figure 9B. In Figure 9C, the percentage of tilted septa was quantified from three independent biological replicates for the indicated strains.

### Microscopy and Image analysis

All imaging was done at room temperature using a Nikon Eclipse Ti-2E microscope with a Yokogawa CSU-WI spinning disk confocal system. This system was equipped with a Photometrics Prime BSI sCMOS camera and a 100x 1.45-NA CFI Plan Apochromat Lambda objective. Cells were grown in YE4S at 25°C unless otherwise stated. All image analysis was performed in ImageJ (Schneider et al., 2012). All statistics and graphs were generated using Prism10 (GraphPad software).

Cell half ratio was determined to quantify division asymmetry in the indicated strains. To do this, the area of each daughter cell was measured using the freehand option in ImageJ. Cell half ratio was determined by dividing the area of the short half of the cell by the area of the long half of the cell (Figure 1B). For all cell half analysis, the cells were stained with Blankophor (MP Biomedicals) to visualize the cell wall and imaged under glass coverslips. Single middle focal plane images were captured. In Figure 1, wild type and phosphatase mutant strains were grown in YE4S at 32°C before being shifted to EMM4S (Edinburgh Minimal Media with 4 Supplements). Cells were maintained in minimal media at 32°C for 48 hours before being imaged. In Figures 2E, 4E, and 8E, cell half ratio was determined from cells grown in YE4S at 25°C. In Figure 2E, the cell half ratio at the initial time of ring assembly was determined using the Rlc1-mNeonGreen signal. In Figure 4I, cells expressing Mid1-mNeonGreen under the control of the endogenous *Pmid1* promoter or *P81nmt1* promoter were grown in EMM4S at 25°C. Cells under the control of the thiamine repressible *P81nmt1* promoter were supplemented with 0.005 μg/ml thiamine for 23 hours. The cell half ratio was determined as described in Figure 1B. In Figure 4I, at least 25 cells were analyzed for each strain.

In Figure 2A, wild type and *par1Δ* strains expressing Rlc1-mNeonGreen were imaged on YE4S agarose pads in 5-minute intervals. Representative single middle focal plane images are shown. The quantifications performed in Figures 2B-2F were determined from time-lapse movies of wild type and *par1Δ* strains expressing Rlc1-mNeonGreen and Sad1-meGFP. Cells were imaged in 5-minute intervals. Z stacks were obtained by acquiring 7 slices with a 0.5µm step size through half the cell. The onset of mitosis was monitored by Sad1-meGFP signal (Nabeshima et al., 1998). In Figure 2B, ring assembly was defined as the time taken to form an intact cytokinetic ring from spindle pole separation. In Figure 2C, ring maturation was defined as the period between cytokinetic ring formation and the onset of constriction. In Figure 2D, ring constriction was defined as the time from the initiation of ring constriction to the disappearance of Rlc1-mNeonGreen from the division site. In Figure 2F, the length from the cell tip to the cytokinetic ring was measured at ring formation. The length was measured again in the same cell at the final moment of cytokinetic ring constriction. Ring displacement was determined as the difference between these two measurements for a cell (Schutt & Moseley, 2020).

To determine fluorescence intensity of wild type and *par1Δ* cells expressing different cytokinetic ring proteins, cells were grown in YE4S at 25°C. Cells were imaged in YE4S and mounted under coverslips on glass slides. Z stacks were obtained by acquiring 13 slices with a 0.5µm step size through the entire cell. To determine the fluorescence intensity of all proteins at the cytokinetic ring, an ROI was drawn around the ring using the rectangular tool in ImageJ. This ROI was used to measure the integrated density from sum intensity projections. The same ROI was moved to a neighboring region without any cells to measure the integrated density. This value was used to subtract background signal. To determine the whole cell fluorescence intensity of Mid1-mNeonGreen, an ROI was drawn around interphase cells in the brightfield image using the freehand option in ImageJ. This ROI was transferred to the corresponding sum projection to measure both the integrated density of the cell and the background signal. In Figure 3G, the fluorescence intensity of Mid1-mNeonGreen in the nucleus was determined in wild type and *par1Δ* cells. These cells also expressed Mid1-mNeonGreen and Nup107-mCherry to mark the nucleus. The cells were imaged on YE4S agarose pads. Nup107-mCherry signal was used to draw an ROI around the nucleus using the freehand option in ImageJ. The same ROI was transferred to a single middle focal plane image and was used to measure the integrated density of Mid1-mNeonGreen. This ROI was also moved to a neighboring area to measure the integrated density of background signal. At least 48 cells were analyzed for each strain in Figure 3G. Welch’s unpaired t-tests were used to compare wild type and *par1Δ* strains in Figure 3.

In Figure 4G-I and Supplemental Figure 3C, cells were grown in EMM4S at 25°C with thiamine to repress the *P81nmt1* promoter (Basi et al., 1993; Maundrell, 1990). Thiamine concentrations ranging from 4 μg/ml to 0.001 μg/ml were tested for controlling *P81nmt1-mid1-mNeonGreen* levels. After 23 hours in 0.005μg/ml thiamine, the *P81nmt1-mid1-mNeonGreen* strain expressed a similar level of Mid1-mNG as we had measured for *par1Δ* cells. Cells were imaged in EMM4S and mounted under coverslips on glass slides. Z stacks were obtained by acquiring 13 slices with a 0.5μm step size through the entire cell. In Figure 4H, the whole cell fluorescence intensity of Mid1-mNeonGreen was determined by drawing an ROI around interphase cells in the brightfield image using the freehand option in ImageJ. This ROI was transferred to the corresponding sum projection to measure both the integrated density of the cell and the background signal. At least 32 cells were analyzed for each strain in Figure 4H. Welch’s unpaired t-tests were used to compare strains in Figure 4H and 4I.

To examine Par1 localization, cells were grown in YE4S at 25°C. These cells expressed Par1-mNeonGreen, Rlc1-mRuby, and Sad1-mCherry. To determine if Par1 and Rlc1 colocalize in Figure 6D, an ROI was drawn around the node region using the rectangular tool in ImageJ. The same ROI was used to determine the plot profile of the selected node region. Par1 and Rlc1 fluorescence were both normalized by setting the minimum fluorescence to 0 and the maximum fluorescence to 100. 9/12 cells had at least partial colocalization between Par1 and Rlc1 at nodes. To determine Par1 localization in the absence of Mid1, wild type and *mid1Δ* cells expressing Par1-mNeonGreen and Rlc1-mRuby were grown in YE4S at 25°C.

### Western blots

Fission yeast whole cell lysates were prepared by growing cells in YE4S to mid-logarithmic phase. Two OD_600_ samples were harvested and rapidly frozen in liquid nitrogen. Cell pellets were resuspended in sample buffer (15% glycerol, 4.5% SDS, 97.5mM Tris pH 6.8, 10% 2-mercaptoethanol, 50mM β-glycerophosphate, 50mM sodium fluoride, 5mM sodium orthovanadate, 1x EDTA-free protease inhibitor cocktail (Sigma Aldrich). The samples were lysed with acid-washed glass beads (Sigma Aldrich) in a Mini-BeadBeater-16 (BioSpec, Bartlesville, OK) for 2 minutes at 4°C. Following this, cell lysates were incubated at 99°C for 5 minutes. Cell lysates were briefly centrifuged, and the supernatant was isolated as the clarified lysate. 5µl of the cell lysates were separated by SDS-PAGE and transferred to a nitrocellulose membrane using the Trans-blot Turbo Transfer System (Bio-Rad). Blots were first probed with anti-myc (SC-40; Santa Cruz) primary antibodies at 4°C. Blots were washed in 1X TBST and then probed with goat anti-mouse secondary antibody (LI-COR). Before being developed on a LiCor Odyssey CLx, blots were washed multiple times in 1X TBST and once in 1X TBS.

To quantify Mid1-13myc levels, a rectangular ROI was drawn around each band in the western blot. The relative band intensities were determined following background subtraction for both Mid1-13myc and the ponceau total protein control. To normalize the values to total protein, the ratio of Mid1-13myc to ponceau were calculated for both wild type and *par1Δ* cells. Fold change was determined by normalizing the Mid1-13myc/ponceau ratio from *par1Δ* to wild type levels. For Figure 3F, the fold change was calculated from three independently grown biological replicates for wild type and *par1Δ* samples. The error bar indicates standard deviation.

### Nuclear Displacement

To displace the nucleus, cells were exponentially grown in YE4S at 25°C. Nuclear displacement experiments were performed as previously described (Almonacid et al., 2009; Daga & Chang, 2005; Opalko et al., 2023). In short, 25µg/ml MBC was added to each culture for 5 minutes at 4°C. Following drug treatment, cells were centrifuged for 8 minutes at 15,000 rpm and resuspended in YE4S with 25µg/ml MBC. Cells were imaged under YE4S+MBC agarose pads. Images were captured in 15-minute intervals. Z stacks were obtained by acquiring 7 slices with a 0.5µm step size through half of the cell.

Maximum intensity projections of wild type and *par1Δ* strains expressing Cut11-mCherry and Rlc1-mNeonGreen following nuclear displacement are shown in Figure 7B. In Supplemental Figure S5, time-lapse movies with wild type and *par1Δ* cells expressing 2xMid1-mNeonGreen, Rlc1-mRuby2, and Cut11-mCherry were analyzed. The time-lapse parameters were the same as described earlier in the nuclear displacement section. The relationship between the position of the cytokinetic ring and displaced nucleus was measured in Figures 7C, S4B, and S6B as follows: the distance from the cell tip closer to the nucleus to the center of the nucleus and ring were measured. To quantify cytokinetic ring and nucleus position, this value was expressed as a fraction of cell length as previously done (Almonacid et al., 2009). For Figures 7C, S5B, and S6B, n ≥ 56 cells were analyzed for each strain.

## ACKNOWLEDGEMENTS

We thank members of the Moseley lab for discussions and comments on the manuscript. This work was supported by a grant from the National Institutes of General Medical Sciences (NIGMS) to J.B.M. (R35GM149248). This work was also supported by bioMT through NIH NIGMS grant P20-GM113132.

## FIGURE LEGENGS

**Figure S1. Additional characterization of *par1Δ* cells.** (A) Representative time-lapse montages of wild type and *par1Δ* cells expressing Rlc1-mNeonGreen and Sad1-mEGFP. Images are a single middle focal plane with inverted fluorescence. Cells were captured in 5-minute intervals. Spindle pole body separation was used as time point 0, which was determined by Sad1-meGFP signal. Scale bar is 7µm. (B) Quantification of Rlc1-mNG recruitment to the cytokinetic ring in wild type and *par1Δ*. Time corresponds to minutes since spindle pole body separation. N ≥ 15 cells each. Errors bars indicate mean and error ± standard deviation. (C) Single middle focal plane images of representative wild type and *par1Δ* cells expressing Cut11-mCherry. Scale bar is 7µm. (D) Quantification of nuclear position. The distance from each cell tip to the center of the nucleus was measured, and then the nuclear position was determined by dividing the short length by the long length. N ≥ 34 cells each. ns indicates *p* value > 0.05 determined by Welch’s unpaired t-test. (E) Time-lapse montage of representative wild type and *par1Δ* cells expressing Cut11-mCherry. Images are a single middle focal plane with mCherry overlaid onto brightfield to visualize the cell tips. Images are captured in 3-minute intervals. Scale bar is 7µm. (F) Quantification of nuclear movement. The distance from the cell tip to the center of the nucleus was measured at every time point. Nuclear movement was determined by averaging the difference between this distance at every time point and dividing it by 3 minutes. N ≥ 15 cells each. ns indicates *p* value > 0.05 determined by Welch’s unpaired t-test.

**Figure S2. Concentration of cytokinesis proteins at the CAR in *par1Δ* cells.** Quantification of the fluorescence intensity for the indicated proteins at the contractile ring (CR) in wild type and par1Δ cells. N ≥ 10 cells each. ns indicates *p* value > 0.05; * indicates *p* value <0.05; ** indicates *p* value < 0.005; *** indicates *p* value <0.0005; **** indicates *p* value <0.0001. All statistical tests performed in this figure are Welch’s unpaired t-tests.

**Figure S3. Mid1 protein levels affect division plane positioning.** (A) Representative images of the indicated strains, mNG images are maximum intensity projections with inverted fluorescence. Scale bar is 14µm. (B) Representative images of cells expressing Mid1-mNG under the control of the thiamine repressible *P81nmt1* promoter. “On” cells were grown with no thiamine for 40 hours (left). “Off” cells were grown in 2.5 μg/ml thiamine (right) for 24 hours. Brightfield images are single middle focal planes. mNG images are sum projections with inverted fluorescence. Scale bar is 12µm. (C) Representative images of the indicated strain. *P81nmt1-mid1-mNG* cells were grown in 0.005μg/ml thiamine for 23 hours. Brightfield images are single middle focal plane. mNG images are sum projections with inverted fluorescence. Scale bar is 12µm.

**Figure S4. Reduced Mid1 levels are specific to PP2A mutants.** (A) Representative images of the indicated strains. Brightfield images are a single middle focal plane. mNG images are maximum intensity projections with inverted fluorescence. Scale bar is 14µm.

**S5. Restoring Mid1 levels in *par1Δ* mutant restores division plane repositioning following nuclear displacement.** (A) Time-lapse montage of wild type and *par1Δ* cells expressing 2xMid1-mNeonGreen, Rlc1-mRuby2, and Cut11-mCherry following MBC treatment and centrifugation. Images are captured in 15-minute intervals. Representative maximum intensity projections are shown with Cut11-mCherry and Rlc1-mRuby2 in magenta and Mid1-mNG in cyan. Brightfield image is a single middle focal plane from the first time point shown. Scale bar is 7µm. (B) Quantification of nuclear displacement experiments. Positions are expressed as a fraction of the cell length. Nuclear positions were separated into <0.3, 0.3-0.35, 0.35-0.40, and >0.4 bins. Error bars correspond to minimum and maximum ring position for each bin. N ≥ 56 cells for each strain.

**Figure S6. Additional examination of Mid1-3xSLiM cells.** (A) Representative images of wild type and *mid1-3xSLiM* cells expressing Mid1-mNeonGreen. Brightfield images are a single middle focal plane. mNG images are maximum intensity projections with inverted fluorescence. Scale bar is 14µm. (B) The relationship between the cytokinetic ring and the displaced nucleus in wild type and *mid1-3xSLIM* cells (as in Figures 7C and S5B). Nuclear positions were separated into <0.3, 0.3-0.35, 0.35-0.40, and >0.4 bins. Error bars correspond to minimum and maximum ring position for each bin. N ≥ 56 cells each. All statistical tests performed in this figure are Welch’s unpaired t-tests.

## REFERENCES

Almonacid, M., Celton-Morizur, S., Jakubowski, J. L., Dingli, F., Loew, D., Mayeux, A., Chen, J.-S., Gould, K. L., Clifford, D. M., & Paoletti, A. (2011). Temporal control of contractile ring assembly by Plo1 regulation of myosin II recruitment by Mid1/anillin. Current Biology: CB, 21(6), 473–479. 10.1016/j.cub.2011.02.003

Almonacid, M., Moseley, J. B., Janvore, J., Mayeux, A., Fraisier, V., Nurse, P., & Paoletti, A. (2009). Spatial control of cytokinesis by Cdr2 kinase and Mid1/anillin nuclear export. Current Biology: CB, 19(11), 961–966. 10.1016/j.cub.2009.04.024

Arasada, R., & Pollard, T. D. (2014). Contractile ring stability in S. pombe depends on F-BAR protein Cdc15p and Bgs1p transport from the Golgi complex. Cell Reports, 8(5), 1533–1544. 10.1016/j.celrep.2014.07.048

Ashraf, S., Tay, Y. D., Kelly, D. A., & Sawin, K. E. (2021). Microtubule-independent movement of the fission yeast nucleus. Journal of Cell Science, 134(6), jcs253021. 10.1242/jcs.253021

Bähler, J., & Pringle, J. R. (1998). Pom1p, a fission yeast protein kinase that provides positional information for both polarized growth and cytokinesis. Genes & Development, 12(9), 1356–1370. 10.1101/gad.12.9.1356

Bähler, J., Steever, A. B., Wheatley, S., Wang, Y. l, Pringle, J. R., Gould, K. L., & McCollum, D. (1998). Role of polo kinase and Mid1p in determining the site of cell division in fission yeast. The Journal of Cell Biology, 143(6), 1603–1616. 10.1083/jcb.143.6.1603

Bähler, J., Wu, J. Q., Longtine, M. S., Shah, N. G., McKenzie, A., Steever, A. B., Wach, A., Philippsen, P., & Pringle, J. R. (1998). Heterologous modules for efficient and versatile PCR-based gene targeting in Schizosaccharomyces pombe. *Yeast (Chichester*, England), 14(10), 943–951. 10.1002/(SICI)1097-0061(199807)14:10<943::AID-YEA292>3.0.CO;2-Y

Balcells, L., Gómez, N., Casamayor, A., Clotet, J., & Ariño, J. (1997). Regulation of salt tolerance in fission yeast by a protein-phosphatase-Z-like Ser/Thr protein phosphatase. European Journal of Biochemistry, 250(2), 476–483. 10.1111/j.1432-1033.1997.0476a.x

Barr, F. A., Elliott, P. R., & Gruneberg, U. (2011). Protein phosphatases and the regulation of mitosis. Journal of Cell Science, 124(Pt 14), 2323–2334. 10.1242/jcs.087106

Barr, F. A., Silljé, H. H. W., & Nigg, E. A. (2004). Polo-like kinases and the orchestration of cell division. Nature Reviews. Molecular Cell Biology, 5(6), 429–440. 10.1038/nrm1401

Basi, G., Schmid, E., & Maundrell, K. (1993). TATA box mutations in the Schizosaccharomyces pombe nmt1 promoter affect transcription efficiency but not the transcription start point or thiamine repressibility. Gene, 123(1), 131–136. 10.1016/0378-1119(93)90552-e

Bastos, R. N., Cundell, M. J., & Barr, F. A. (2014). KIF4A and PP2A-B56 form a spatially restricted feedback loop opposing Aurora B at the anaphase central spindle. The Journal of Cell Biology, 207(6), 683–693. 10.1083/jcb.201409129

Bhavsar-Jog, Y. P., & Bi, E. (2017). Mechanics and regulation of cytokinesis in budding yeast. Seminars in Cell & Developmental Biology, 66, 107–118. 10.1016/j.semcdb.2016.12.010

Bimbó, A., Liu, J., & Balasubramanian, M. K. (2005). Roles of Pdk1p, a fission yeast protein related to phosphoinositide-dependent protein kinase, in the regulation of mitosis and cytokinesis. Molecular Biology of the Cell, 16(7), 3162–3175. 10.1091/mbc.e04-09-0769

Bohnert, K. A., & Gould, K. L. (2011). On the cutting edge: Post-translational modifications in cytokinesis. Trends in Cell Biology, 21(5), 283–292. 10.1016/j.tcb.2011.01.006

Bollen, M., Gerlich, D. W., & Lesage, B. (2009). Mitotic phosphatases: From entry guards to exit guides. Trends in Cell Biology, 19(10), 531–541. 10.1016/j.tcb.2009.06.005

Brennan, I. M., Peters, U., Kapoor, T. M., & Straight, A. F. (2007). Polo-like kinase controls vertebrate spindle elongation and cytokinesis. PloS One, 2(5), e409. 10.1371/journal.pone.0000409

Burgess, A., Vigneron, S., Brioudes, E., Labbé, J.-C., Lorca, T., & Castro, A. (2010). Loss of human Greatwall results in G2 arrest and multiple mitotic defects due to deregulation of the cyclin B-Cdc2/PP2A balance. Proceedings of the National Academy of Sciences of the United States of America, 107(28), 12564–12569. 10.1073/pnas.0914191107

Burkard, M. E., Maciejowski, J., Rodriguez-Bravo, V., Repka, M., Lowery, D. M., Clauser, K. R., Zhang, C., Shokat, K. M., Carr, S. A., Yaffe, M. B., & Jallepalli, P. V. (2009). Plk1 self-organization and priming phosphorylation of HsCYK-4 at the spindle midzone regulate the onset of division in human cells. PLoS Biology, 7(5), e1000111. 10.1371/journal.pbio.1000111

Carmena, M., Wheelock, M., Funabiki, H., & Earnshaw, W. C. (2012). The chromosomal passenger complex (CPC): From easy rider to the godfather of mitosis. Nature Reviews. Molecular Cell Biology, 13(12), 789–803. 10.1038/nrm3474

Celton-Morizur, S., Bordes, N., Fraisier, V., Tran, P. T., & Paoletti, A. (2004). C-terminal anchoring of mid1p to membranes stabilizes cytokinetic ring position in early mitosis in fission yeast. Molecular and Cellular Biology, 24(24), 10621–10635. 10.1128/MCB.24.24.10621-10635.2004

Chang, F., Woollard, A., & Nurse, P. (1996). Isolation and characterization of fission yeast mutants defective in the assembly and placement of the contractile actin ring. Journal of Cell Science, 109 ( *Pt 1*), 131–142. 10.1242/jcs.109.1.131

Cheffings, T. H., Burroughs, N. J., & Balasubramanian, M. K. (2016). Actomyosin Ring Formation and Tension Generation in Eukaryotic Cytokinesis. Current Biology: CB, 26(15), R719–R737. 10.1016/j.cub.2016.06.071

Clifford, D. M., Wolfe, B. A., Roberts-Galbraith, R. H., McDonald, W. H., Yates, J. R., & Gould, K. L. (2008). The Clp1/Cdc14 phosphatase contributes to the robustness of cytokinesis by association with anillin-related Mid1. The Journal of Cell Biology, 181(1), 79–88. 10.1083/jcb.200709060

Cundell, M. J., Hutter, L. H., Nunes Bastos, R., Poser, E., Holder, J., Mohammed, S., Novak, B., & Barr, F. A. (2016). A PP2A-B55 recognition signal controls substrate dephosphorylation kinetics during mitotic exit. The Journal of Cell Biology, 214(5), 539–554. 10.1083/jcb.201606033

Daga, R. R., & Chang, F. (2005). Dynamic positioning of the fission yeast cell division plane. Proceedings of the National Academy of Sciences of the United States of America, 102(23), 8228–8232. 10.1073/pnas.0409021102

Dai, X., Chen, X., Hakizimana, O., & Mei, Y. (2019). Genetic interactions between ANLN and KDR are prognostic for breast cancer survival. Oncology Reports, 42(6), 2255–2266. 10.3892/or.2019.7332

D’Avino, P. P., Giansanti, M. G., & Petronczki, M. (2015). Cytokinesis in animal cells. Cold Spring Harbor Perspectives in Biology, 7(4), a015834. 10.1101/cshperspect.a015834

Echard, A., Hickson, G. R. X., Foley, E., & O’Farrell, P. H. (2004). Terminal cytokinesis events uncovered after an RNAi screen. Current Biology: CB, 14(18), 1685– 1693. 10.1016/j.cub.2004.08.063

Eng, K., Naqvi, N. I., Wong, K. C., & Balasubramanian, M. K. (1998). Rng2p, a protein required for cytokinesis in fission yeast, is a component of the actomyosin ring and the spindle pole body. Current Biology: CB, 8(11), 611–621. 10.1016/s0960-9822(98)70248-9

García-Blanco, N., Vázquez-Bolado, A., & Moreno, S. (2019). Greatwall-Endosulfine: A Molecular Switch that Regulates PP2A/B55 Protein Phosphatase Activity in Dividing and Quiescent Cells. International Journal of Molecular Sciences, 20(24), 6228. 10.3390/ijms20246228

Goyal, A., & Simanis, V. (2012). Characterization of ypa1 and ypa2, the Schizosaccharomyces pombe orthologs of the peptidyl proyl isomerases that activate PP2A, reveals a role for Ypa2p in the regulation of cytokinesis. Genetics, 190(4), 1235–1250. 10.1534/genetics.111.138040

Green, R. A., Paluch, E., & Oegema, K. (2012). Cytokinesis in animal cells. Annual Review of Cell and Developmental Biology, 28, 29–58. 10.1146/annurev-cellbio-101011-155718

Guzman-Vendrell, M., Baldissard, S., Almonacid, M., Mayeux, A., Paoletti, A., & Moseley, J. B. (2013). Blt1 and Mid1 provide overlapping membrane anchors to position the division plane in fission yeast. Molecular and Cellular Biology, 33(2), 418–428. 10.1128/MCB.01286-12

Hall, P. A., Todd, C. B., Hyland, P. L., McDade, S. S., Grabsch, H., Dattani, M., Hillan, K. J., & Russell, S. E. H. (2005). The septin-binding protein anillin is overexpressed in diverse human tumors. Clinical Cancer Research: An Official Journal of the American Association for Cancer Research, *11*(19 Pt 1), 6780– 6786. 10.1158/1078-0432.CCR-05-0997

Healy, A. M., Zolnierowicz, S., Stapleton, A. E., Goebl, M., DePaoli-Roach, A. A., & Pringle, J. R. (1991). CDC55, a Saccharomyces cerevisiae gene involved in cellular morphogenesis: Identification, characterization, and homology to the B subunit of mammalian type 2A protein phosphatase. Molecular and Cellular Biology, 11(11), 5767–5780. 10.1128/mcb.11.11.5767-5780.1991

Henson, J. H., Ditzler, C. E., Germain, A., Irwin, P. M., Vogt, E. T., Yang, S., Wu, X., & Shuster, C. B. (2017). The ultrastructural organization of actin and myosin II filaments in the contractile ring: New support for an old model of cytokinesis. Molecular Biology of the Cell, 28(5), 613–623. 10.1091/mbc.E16-06-0466

Hertz, E. P. T., Kruse, T., Davey, N. E., López-Méndez, B., Sigurðsson, J. O., Montoya, G., Olsen, J. V., & Nilsson, J. (2016). A Conserved Motif Provides Binding Specificity to the PP2A-B56 Phosphatase. Molecular Cell, 63(4), 686–695. 10.1016/j.molcel.2016.06.024

Hickson, G. R. X., & O’Farrell, P. H. (2008). Rho-dependent control of anillin behavior during cytokinesis. The Journal of Cell Biology, 180(2), 285–294. 10.1083/jcb.200709005

Holder, J., Poser, E., & Barr, F. A. (2019). Getting out of mitosis: Spatial and temporal control of mitotic exit and cytokinesis by PP1 and PP2A. FEBS Letters, 593(20), 2908–2924. 10.1002/1873-3468.13595

Janssens, V., & Goris, J. (2001). Protein phosphatase 2A: A highly regulated family of serine/threonine phosphatases implicated in cell growth and signalling. The Biochemical Journal, 353(Pt 3), 417–439. 10.1042/0264-6021:3530417

Jiang, W., & Hallberg, R. L. (2000). Isolation and characterization of par1(+) and par2(+): Two Schizosaccharomyces pombe genes encoding B’ subunits of protein phosphatase 2A. Genetics, 154(3), 1025–1038. 10.1093/genetics/154.3.1025

Jiang, W., & Hallberg, R. L. (2001). Correct regulation of the septation initiation network in Schizosaccharomyces pombe requires the activities of par1 and par2. Genetics, 158(4), 1413–1429. 10.1093/genetics/158.4.1413

Kim, H., Johnson, J. M., Lera, R. F., Brahma, S., & Burkard, M. E. (2017). Anillin Phosphorylation Controls Timely Membrane Association and Successful Cytokinesis. PLoS Genetics, 13(1), e1006511. 10.1371/journal.pgen.1006511

Kinoshita, K., Nemoto, T., Nabeshima, K., Kondoh, H., Niwa, H., & Yanagida, M. (1996). The regulatory subunits of fission yeast protein phosphatase 2A (PP2A) affect cell morphogenesis, cell wall synthesis and cytokinesis. Genes to Cells: Devoted to Molecular & Cellular Mechanisms, 1(1), 29–45. 10.1046/j.1365-2443.1996.02002.x

Kinoshita, N., Yamano, H., Niwa, H., Yoshida, T., & Yanagida, M. (1993). Negative regulation of mitosis by the fission yeast protein phosphatase ppa2. Genes & Development, 7(6), 1059–1071. 10.1101/gad.7.6.1059

Kruse, T., Gnosa, S. P., Nasa, I., Garvanska, D. H., Hein, J. B., Nguyen, H., Samsøe-Petersen, J., Lopez-Mendez, B., Hertz, E. P. T., Schwarz, J., Pena, H. S., Nikodemus, D., Kveiborg, M., Kettenbach, A. N., & Nilsson, J. (2020). Mechanisms of site-specific dephosphorylation and kinase opposition imposed by PP2A regulatory subunits. The EMBO Journal, 39(13), e103695. 10.15252/embj.2019103695

Le Goff, X., Buvelot, S., Salimova, E., Guerry, F., Schmidt, S., Cueille, N., Cano, E., & Simanis, V. (2001). The protein phosphatase 2A B’-regulatory subunit par1p is implicated in regulation of the S. pombe septation initiation network. FEBS Letters, 508(1), 136–142. 10.1016/s0014-5793(01)03047-2

Magliozzi, J. O., & Moseley, J. B. (2021). Connecting cell polarity signals to the cytokinetic machinery in yeast and metazoan cells. *Cell Cycle (Georgetown*, Tex.), 20(1), 1–10. 10.1080/15384101.2020.1864941

Magliozzi, J. O., Sears, J., Cressey, L., Brady, M., Opalko, H. E., Kettenbach, A. N., & Moseley, J. B. (2020). Fission yeast Pak1 phosphorylates anillin-like Mid1 for spatial control of cytokinesis. The Journal of Cell Biology, 219(8), e201908017. 10.1083/jcb.201908017

Magnusson, K., Gremel, G., Rydén, L., Pontén, V., Uhlén, M., Dimberg, A., Jirström, K., & Pontén, F. (2016). ANLN is a prognostic biomarker independent of Ki-67 and essential for cell cycle progression in primary breast cancer. BMC Cancer, 16(1), 904. 10.1186/s12885-016-2923-8

Mangione, M. C., & Gould, K. L. (2019). Molecular form and function of the cytokinetic ring. Journal of Cell Science, 132(12), jcs226928. 10.1242/jcs.226928

Martín-García, R., Arribas, V., Coll, P. M., Pinar, M., Viana, R. A., Rincón, S. A., Correa-Bordes, J., Ribas, J. C., & Pérez, P. (2018). Paxillin-Mediated Recruitment of Calcineurin to the Contractile Ring Is Required for the Correct Progression of Cytokinesis in Fission Yeast. Cell Reports, 25(3), 772–783.e4. 10.1016/j.celrep.2018.09.062

Martín-García, R., Coll, P. M., & Pérez, P. (2014). F-BAR domain protein Rga7 collaborates with Cdc15 and Imp2 to ensure proper cytokinesis in fission yeast. Journal of Cell Science, 127(Pt 19), 4146–4158. 10.1242/jcs.146233

Maundrell, K. (1990). nmt1 of fission yeast. A highly transcribed gene completely repressed by thiamine. The Journal of Biological Chemistry, 265(19), 10857–10864.

McDonald, N. A., Vander Kooi, C. W., Ohi, M. D., & Gould, K. L. (2015). Oligomerization but Not Membrane Bending Underlies the Function of Certain F-BAR Proteins in Cell Motility and Cytokinesis. Developmental Cell, 35(6), 725–736. 10.1016/j.devcel.2015.11.023

Mishra, M., Huang, Y., Srivastava, P., Srinivasan, R., Sevugan, M., Shlomovitz, R., Gov, N., Rao, M., & Balasubramanian, M. (2012). Cylindrical cellular geometry ensures fidelity of division site placement in fission yeast. Journal of Cell Science, 125(Pt 16), 3850–3857. 10.1242/jcs.103788

Mishra, M., Karagiannis, J., Trautmann, S., Wang, H., McCollum, D., & Balasubramanian, M. K. (2004). The Clp1p/Flp1p phosphatase ensures completion of cytokinesis in response to minor perturbation of the cell division machinery in Schizosaccharomyces pombe. Journal of Cell Science, 117(Pt 17), 3897–3910. 10.1242/jcs.01244

Moreno, S., Klar, A., & Nurse, P. (1991). Molecular genetic analysis of fission yeast Schizosaccharomyces pombe. Methods in Enzymology, 194, 795–823. 10.1016/0076-6879(91)94059-l

Motegi, F., Mishra, M., Balasubramanian, M. K., & Mabuchi, I. (2004). Myosin-II reorganization during mitosis is controlled temporally by its dephosphorylation and spatially by Mid1 in fission yeast. The Journal of Cell Biology, 165(5), 685– 695. 10.1083/jcb.200402097

Moyano-Rodriguez, Y., & Queralt, E. (2019). PP2A Functions during Mitosis and Cytokinesis in Yeasts. International Journal of Molecular Sciences, 21(1), 264. 10.3390/ijms21010264

Moyano-Rodríguez, Y., Vaquero, D., Vilalta-Castany, O., Foltman, M., Sanchez-Diaz, A., & Queralt, E. (2022). PP2A-Cdc55 phosphatase regulates actomyosin ring contraction and septum formation during cytokinesis. Cellular and Molecular Life Sciences: CMLS, 79(3), 165. 10.1007/s00018-022-04209-1

Nabeshima, K., Nakagawa, T., Straight, A. F., Murray, A., Chikashige, Y., Yamashita, Y. M., Hiraoka, Y., & Yanagida, M. (1998). Dynamics of centromeres during metaphase-anaphase transition in fission yeast: Dis1 is implicated in force balance in metaphase bipolar spindle. Molecular Biology of the Cell, 9(11), 3211– 3225. 10.1091/mbc.9.11.3211

Nilsson, J. (2019). Protein phosphatases in the regulation of mitosis. The Journal of Cell Biology, 218(2), 395–409. 10.1083/jcb.201809138

Opalko, H., Geng, S., Hall, A. R., Vavylonis, D., & Moseley, J. B. (2023). Design principles of Cdr2 node patterns in fission yeast cells. Molecular Biology of the Cell, 34(11), br18. 10.1091/mbc.E23-04-0135

Padmanabhan, A., Bakka, K., Sevugan, M., Naqvi, N. I., D’souza, V., Tang, X., Mishra, M., & Balasubramanian, M. K. (2011). IQGAP-related Rng2p organizes cortical nodes and ensures position of cell division in fission yeast. Current Biology: CB, 21(6), 467–472. 10.1016/j.cub.2011.01.059

Paoletti, A., & Chang, F. (2000). Analysis of mid1p, a protein required for placement of the cell division site, reveals a link between the nucleus and the cell surface in fission yeast. Molecular Biology of the Cell, 11(8), 2757–2773. 10.1091/mbc.11.8.2757

Peris, I., Romero-Murillo, S., Vicente, C., Narla, G., & Odero, M. D. (2023). Regulation and role of the PP2A-B56 holoenzyme family in cancer. Biochimica Et Biophysica Acta. Reviews on Cancer, 1878(5), 188953. 10.1016/j.bbcan.2023.188953

Piekny, A. J., & Glotzer, M. (2008). Anillin is a scaffold protein that links RhoA, actin, and myosin during cytokinesis. Current Biology: CB, 18(1), 30–36. 10.1016/j.cub.2007.11.068

Pollard, T. D., & O’Shaughnessy, B. (2019). Molecular Mechanism of Cytokinesis. Annual Review of Biochemistry, 88, 661–689. 10.1146/annurev-biochem-062917-012530

Pollard, T. D., & Wu, J.-Q. (2010). Understanding cytokinesis: Lessons from fission yeast. Nature Reviews. Molecular Cell Biology, 11(2), 149–155. 10.1038/nrm2834

Proctor, S. A., Minc, N., Boudaoud, A., & Chang, F. (2012). Contributions of turgor pressure, the contractile ring, and septum assembly to forces in cytokinesis in fission yeast. Current Biology: CB, 22(17), 1601–1608. 10.1016/j.cub.2012.06.042

Rezig, I. M., Yaduma, W. G., Gould, G. W., & McInerny, C. J. (2021). Anillin/Mid1p interacts with the ESCRT-associated protein Vps4p and mitotic kinases to regulate cytokinesis in fission yeast. *Cell Cycle (Georgetown*, Tex.), 20(18), 1845–1860. 10.1080/15384101.2021.1962637

Rezig, I. M., Yaduma, W. G., Gould, G. W., & McInerny, C. J. (2023). The role of anillin/Mid1p during medial division and cytokinesis: From fission yeast to cancer cells. *Cell Cycle (Georgetown*, Tex.), 22(6), 633–644. 10.1080/15384101.2022.2147655

Rincon, S. A., & Paoletti, A. (2012). Mid1/anillin and the spatial regulation of cytokinesis in fission yeast. Cytoskeleton (Hoboken, N.J.), 69(10), 764–777. 10.1002/cm.21056

Rincon, S. A., & Paoletti, A. (2016). Molecular control of fission yeast cytokinesis. Seminars in Cell & Developmental Biology, 53, 28–38. 10.1016/j.semcdb.2016.01.007

Roberts-Galbraith, R. H., Chen, J.-S., Wang, J., & Gould, K. L. (2009). The SH3 domains of two PCH family members cooperate in assembly of the Schizosaccharomyces pombe contractile ring. The Journal of Cell Biology, 184(1), 113–127. 10.1083/jcb.200806044

Santamaria, A., Neef, R., Eberspächer, U., Eis, K., Husemann, M., Mumberg, D., Prechtl, S., Schulze, V., Siemeister, G., Wortmann, L., Barr, F. A., & Nigg, E. A. (2007). Use of the novel Plk1 inhibitor ZK-thiazolidinone to elucidate functions of Plk1 in early and late stages of mitosis. Molecular Biology of the Cell, 18(10), 4024–4036. 10.1091/mbc.e07-05-0517

Schneider, C. A., Rasband, W. S., & Eliceiri, K. W. (2012). NIH Image to ImageJ: 25 years of image analysis. Nature Methods, 9(7), 671–675. 10.1038/nmeth.2089

Schutt, K. L., & Moseley, J. B. (2020). The phosphatase inhibitor Sds23 promotes symmetric spindle positioning in fission yeast. *Cytoskeleton (Hoboken*, N.J.), 77(12), 544–557. 10.1002/cm.21648

Sechi, S., Piergentili, R., & Giansanti, M. G. (2022). Minor Kinases with Major Roles in Cytokinesis Regulation. Cells, 11(22), 3639. 10.3390/cells11223639

Seshacharyulu, P., Pandey, P., Datta, K., & Batra, S. K. (2013). Phosphatase: PP2A structural importance, regulation and its aberrant expression in cancer. Cancer Letters, 335(1), 9–18. 10.1016/j.canlet.2013.02.036

Snider, C. E., Chandra, M., McDonald, N. A., Willet, A. H., Collier, S. E., Ohi, M. D., Jackson, L. P., & Gould, K. L. (2020). Opposite Surfaces of the Cdc15 F-BAR Domain Create a Membrane Platform That Coordinates Cytoskeletal and Signaling Components for Cytokinesis. Cell Reports, 33(12), 108526. 10.1016/j.celrep.2020.108526

Sohrmann, M., Fankhauser, C., Brodbeck, C., & Simanis, V. (1996). The dmf1/mid1 gene is essential for correct positioning of the division septum in fission yeast. Genes & Development, 10(21), 2707–2719. 10.1101/gad.10.21.2707

Somma, M. P., Fasulo, B., Cenci, G., Cundari, E., & Gatti, M. (2002). Molecular dissection of cytokinesis by RNA interference in Drosophila cultured cells. Molecular Biology of the Cell, 13(7), 2448–2460. 10.1091/mbc.01-12-0589

Straight, A. F., Field, C. M., & Mitchison, T. J. (2005). Anillin binds nonmuscle myosin II and regulates the contractile ring. Molecular Biology of the Cell, 16(1), 193–201. 10.1091/mbc.e04-08-0758

Su, K.-C., Takaki, T., & Petronczki, M. (2011). Targeting of the RhoGEF Ect2 to the equatorial membrane controls cleavage furrow formation during cytokinesis. Developmental Cell, 21(6), 1104–1115. 10.1016/j.devcel.2011.11.003

Suijkerbuijk, S. J. E., Vleugel, M., Teixeira, A., & Kops, G. J. P. L. (2012). Integration of kinase and phosphatase activities by BUBR1 ensures formation of stable kinetochore-microtubule attachments. Developmental Cell, 23(4), 745–755. 10.1016/j.devcel.2012.09.005

Tolic-Nørrelykke, I. M., Sacconi, L., Stringari, C., Raabe, I., & Pavone, F. S. (2005). Nuclear and division-plane positioning revealed by optical micromanipulation. Current Biology: CB, 15(13), 1212–1216. 10.1016/j.cub.2005.05.052

Trautmann, S., Wolfe, B. A., Jorgensen, P., Tyers, M., Gould, K. L., & McCollum, D. (2001). Fission yeast Clp1p phosphatase regulates G2/M transition and coordination of cytokinesis with cell cycle progression. Current Biology: CB, 11(12), 931–940. 10.1016/s0960-9822(01)00268-8

Trinkle-Mulcahy, L., & Lamond, A. I. (2006). Mitotic phosphatases: No longer silent partners. Current Opinion in Cell Biology, 18(6), 623–631. 10.1016/j.ceb.2006.09.001

Uhlén, M., Fagerberg, L., Hallström, B. M., Lindskog, C., Oksvold, P., Mardinoglu, A., Sivertsson, Å., Kampf, C., Sjöstedt, E., Asplund, A., Olsson, I., Edlund, K., Lundberg, E., Navani, S., Szigyarto, C. A.-K., Odeberg, J., Djureinovic, D., Takanen, J. O., Hober, S., … Pontén, F. (2015). Proteomics. Tissue-based map of the human proteome. Science (New York, N.Y.), 347(6220), 1260419. 10.1126/science.1260419

Vallardi, G., Allan, L. A., Crozier, L., & Saurin, A. T. (2019). Division of labour between PP2A-B56 isoforms at the centromere and kinetochore. eLife, 8, e42619. 10.7554/eLife.42619

van der Waal, M. S., Hengeveld, R. C. C., van der Horst, A., & Lens, S. M. A. (2012). Cell division control by the Chromosomal Passenger Complex. Experimental Cell Research, 318(12), 1407–1420. 10.1016/j.yexcr.2012.03.015

Wang, D., Naydenov, N. G., Dozmorov, M. G., Koblinski, J. E., & Ivanov, A. I. (2020). Anillin regulates breast cancer cell migration, growth, and metastasis by non-canonical mechanisms involving control of cell stemness and differentiation. Breast Cancer Research: BCR, 22(1), 3. 10.1186/s13058-019-1241-x

Wang, J., Wang, Z., Yu, T., Yang, H., Virshup, D. M., Kops, G. J. P. L., Lee, S. H., Zhou, W., Li, X., Xu, W., & Rao, Z. (2016). Crystal structure of a PP2A B56-BubR1 complex and its implications for PP2A substrate recruitment and localization. Protein & Cell, 7(7), 516–526. 10.1007/s13238-016-0283-4

Wang, X., Bajaj, R., Bollen, M., Peti, W., & Page, R. (2016). Expanding the PP2A Interactome by Defining a B56-Specific SLiM. Structure (London, England: 1993), 24(12), 2174–2181. 10.1016/j.str.2016.09.010

Wang, Z., Chen, J., Zhong, M.-Z., Huang, J., Hu, Y.-P., Feng, D.-Y., Zhou, Z.-J., Luo, X., Liu, Z.-Q., Jiang, W.-Z., & Zhou, W.-B. (2017). Overexpression of ANLN contributed to poor prognosis of anthracycline-based chemotherapy in breast cancer patients. Cancer Chemotherapy and Pharmacology, 79(3), 535–543. 10.1007/s00280-017-3248-2

Weaver, B. A. A., Silk, A. D., Montagna, C., Verdier-Pinard, P., & Cleveland, D. W. (2007). Aneuploidy acts both oncogenically and as a tumor suppressor. Cancer Cell, 11(1), 25–36. 10.1016/j.ccr.2006.12.003

Willet, A. H., DeWitt, A. K., Beckley, J. R., Clifford, D. M., & Gould, K. L. (2019a). NDR Kinase Sid2 Drives Anillin-like Mid1 from the Membrane to Promote Cytokinesis and Medial Division Site Placement. Current Biology: CB, 29(6), 1055–1063.e2. 10.1016/j.cub.2019.01.075

Willet, A. H., DeWitt, A. K., Beckley, J. R., Clifford, D. M., & Gould, K. L. (2019b). NDR Kinase Sid2 Drives Anillin-like Mid1 from the Membrane to Promote Cytokinesis and Medial Division Site Placement. Current Biology: CB, 29(6), 1055–1063.e2. 10.1016/j.cub.2019.01.075

Wolfe, B. A., Takaki, T., Petronczki, M., & Glotzer, M. (2009). Polo-like kinase 1 directs assembly of the HsCyk-4 RhoGAP/Ect2 RhoGEF complex to initiate cleavage furrow formation. PLoS Biology, 7(5), e1000110. 10.1371/journal.pbio.1000110

Wu, C.-G., Chen, H., Guo, F., Yadav, V. K., Mcilwain, S. J., Rowse, M., Choudhary, A., Lin, Z., Li, Y., Gu, T., Zheng, A., Xu, Q., Lee, W., Resch, E., Johnson, B., Day, J., Ge, Y., Ong, I. M., Burkard, M. E., … Xing, Y. (2017). PP2A-B’ holoenzyme substrate recognition, regulation and role in cytokinesis. Cell Discovery, 3, 17027. 10.1038/celldisc.2017.27

Wu, J.-Q., Kuhn, J. R., Kovar, D. R., & Pollard, T. D. (2003). Spatial and temporal pathway for assembly and constriction of the contractile ring in fission yeast cytokinesis. Developmental Cell, 5(5), 723–734. 10.1016/s1534-5807(03)00324-1

Wu, J.-Q., Sirotkin, V., Kovar, D. R., Lord, M., Beltzner, C. C., Kuhn, J. R., & Pollard, T. D. (2006). Assembly of the cytokinetic contractile ring from a broad band of nodes in fission yeast. The Journal of Cell Biology, 174(3), 391–402. 10.1083/jcb.200602032

Yoshida, T., Toda, T., & Yanagida, M. (1994). A calcineurin-like gene ppb1+ in fission yeast: Mutant defects in cytokinesis, cell polarity, mating and spindle pole body positioning. Journal of Cell Science, 107 ( *Pt 7*), 1725–1735. 10.1242/jcs.107.7.1725

Zhao, W.-M., & Fang, G. (2005). Anillin is a substrate of anaphase-promoting complex/cyclosome (APC/C) that controls spatial contractility of myosin during late cytokinesis. The Journal of Biological Chemistry, 280(39), 33516–33524. 10.1074/jbc.M504657200

Zhou, W., Wang, Z., Shen, N., Pi, W., Jiang, W., Huang, J., Hu, Y., Li, X., & Sun, L. (2015). Knockdown of ANLN by lentivirus inhibits cell growth and migration in human breast cancer. Molecular and Cellular Biochemistry, 398(1–2), 11–19. 10.1007/s11010-014-2200-6

