## Supplementary figures and images for "PP2A-B56 regulates Mid1 protein levels for proper cytokinesis in fission yeast"

### Supplemental Figures S1-S6

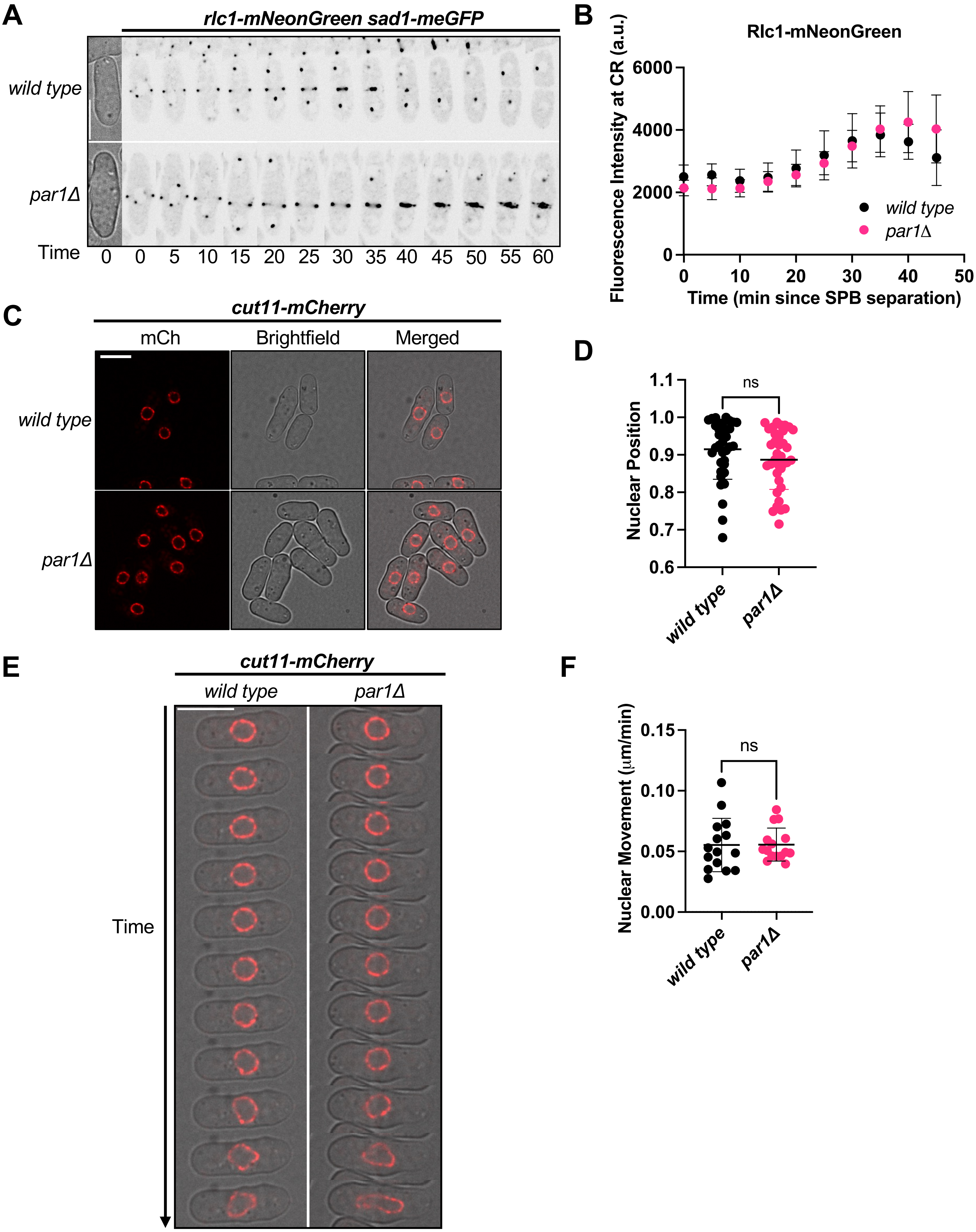

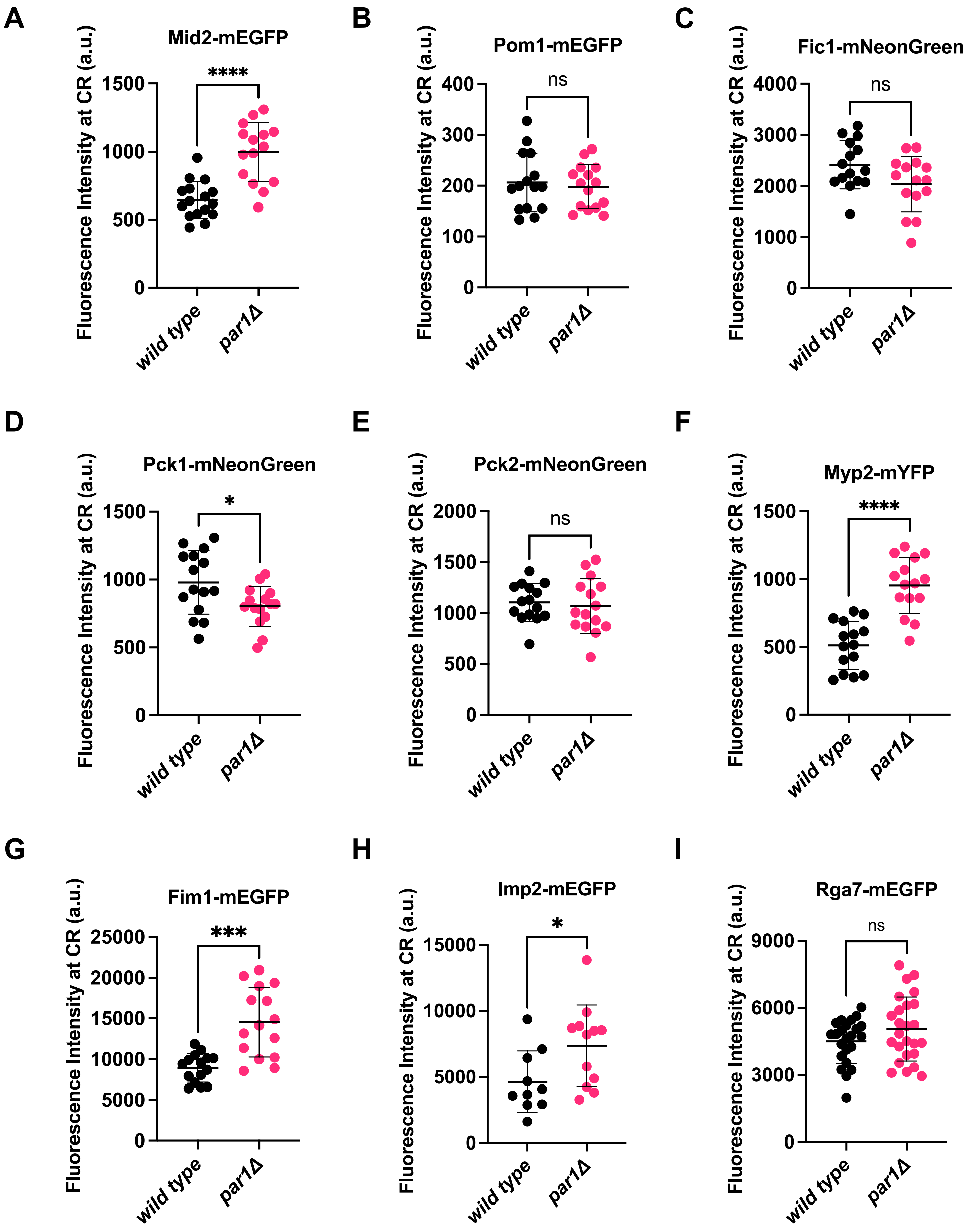

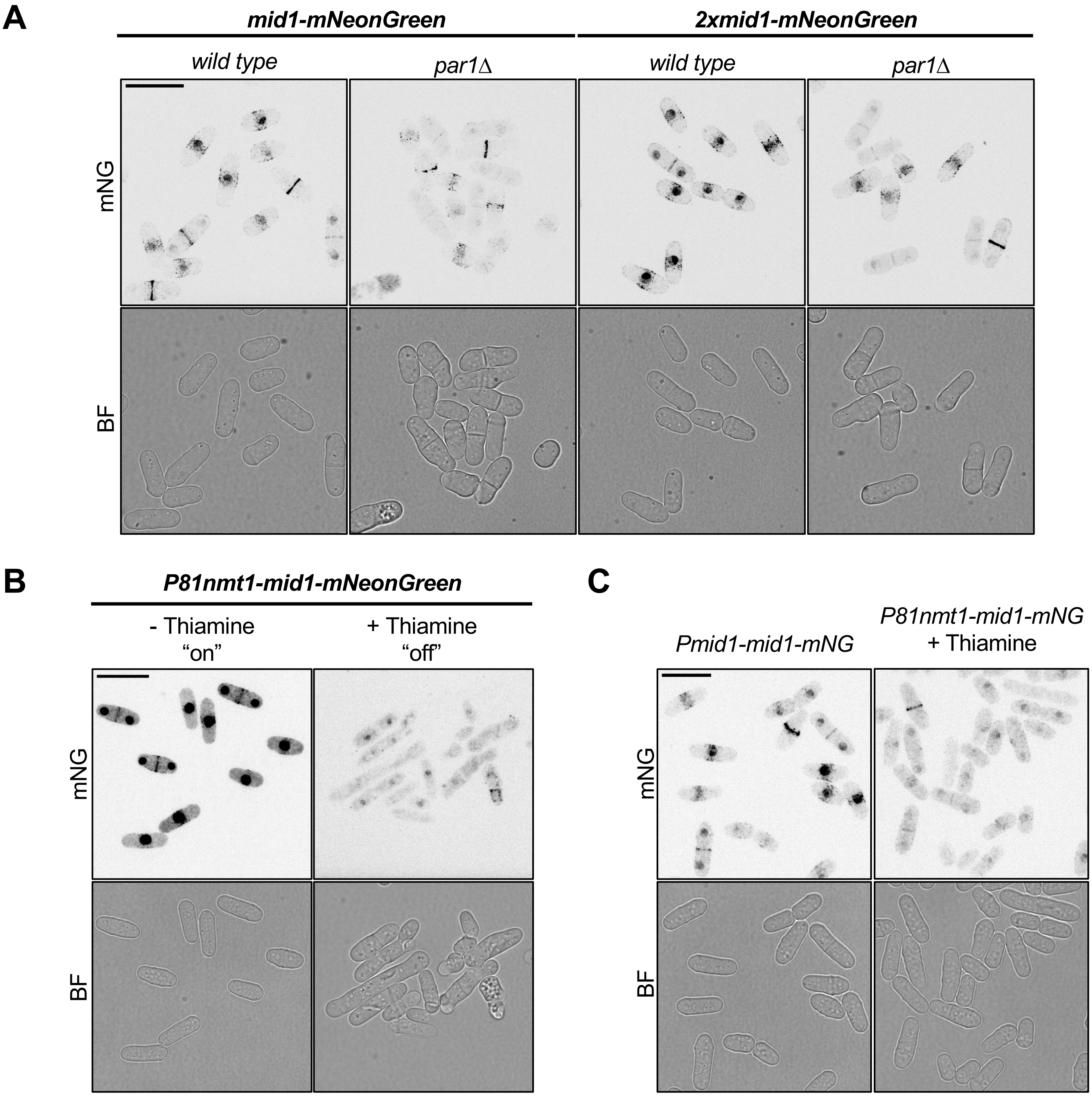

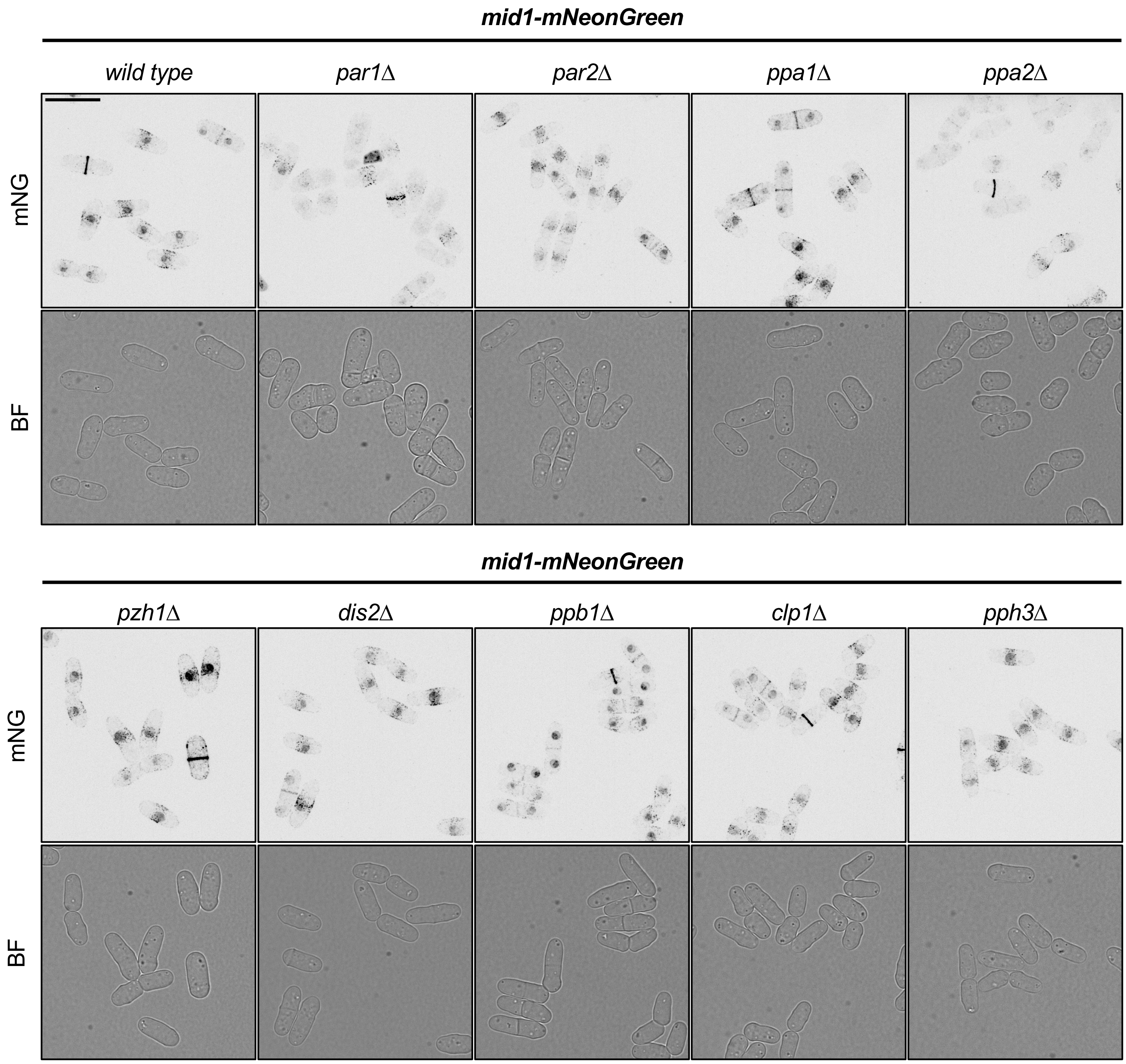

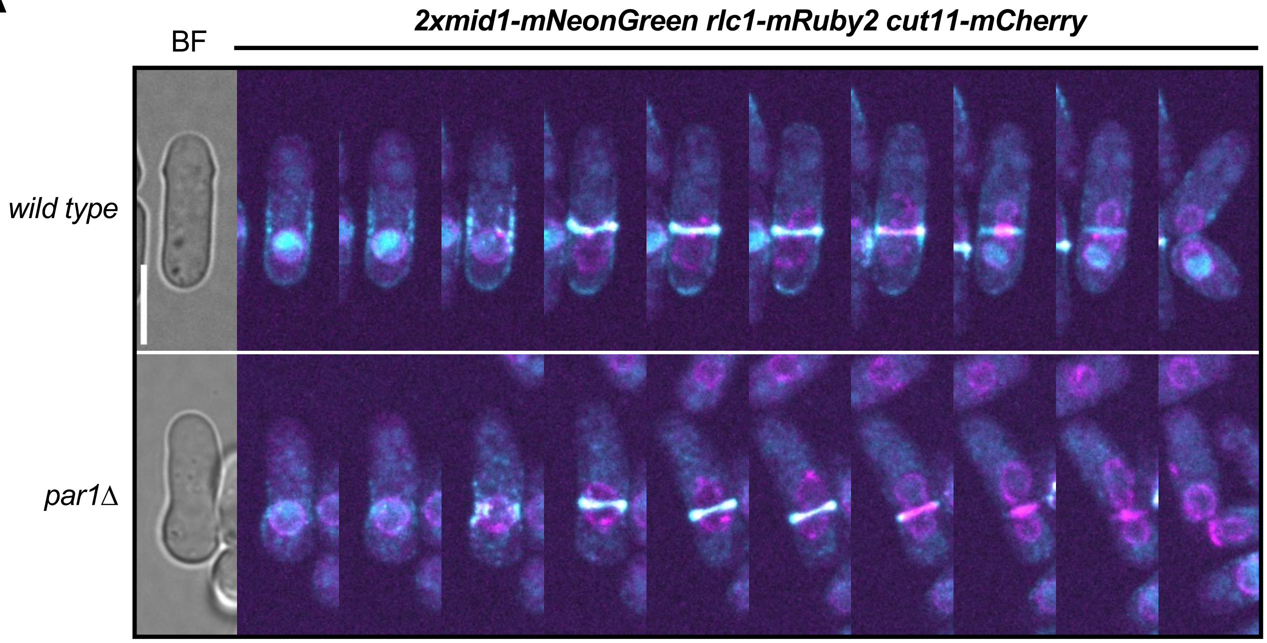

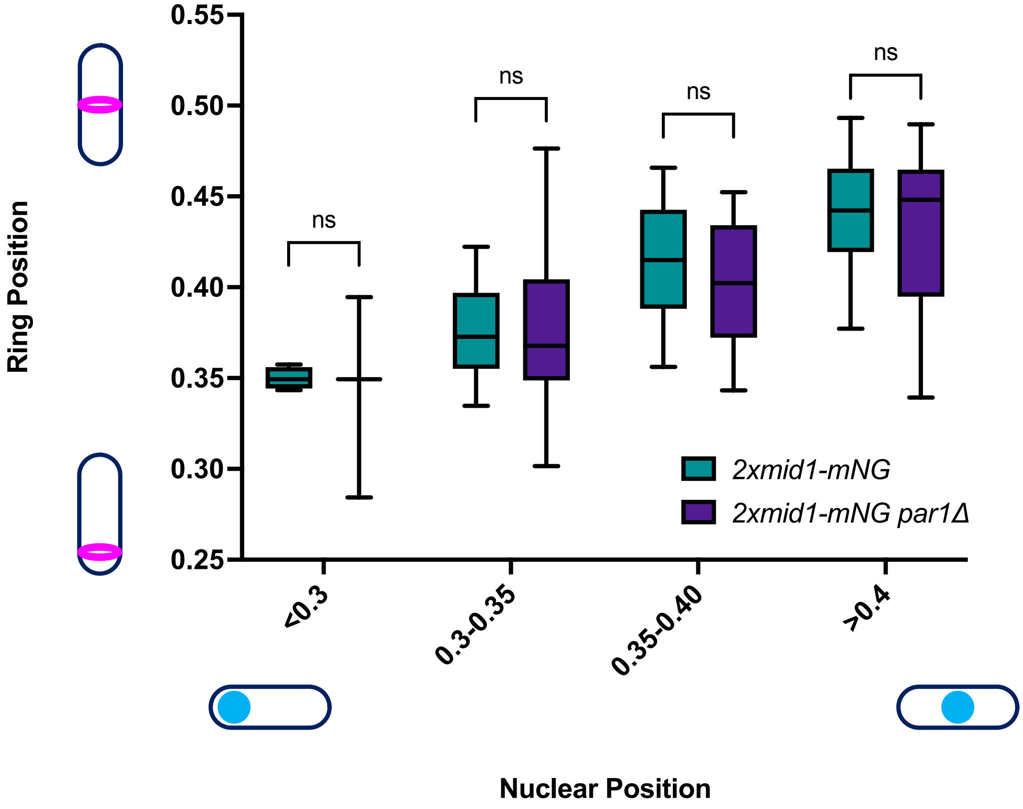

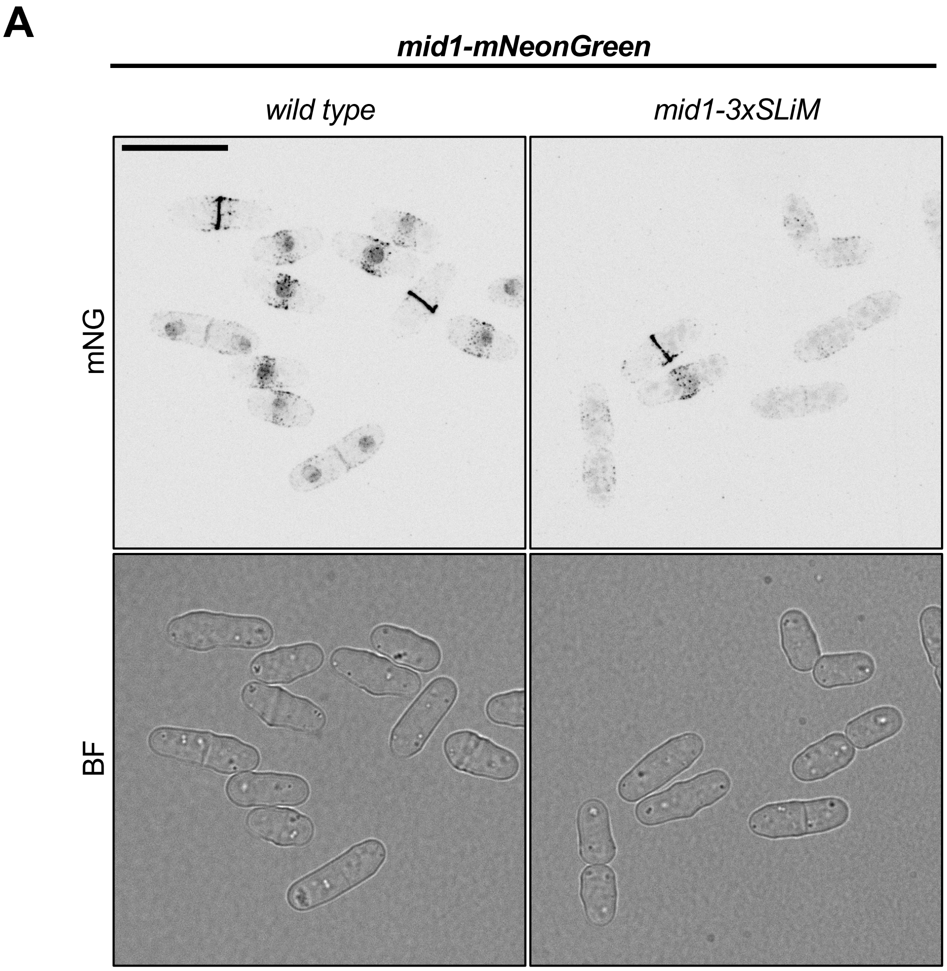

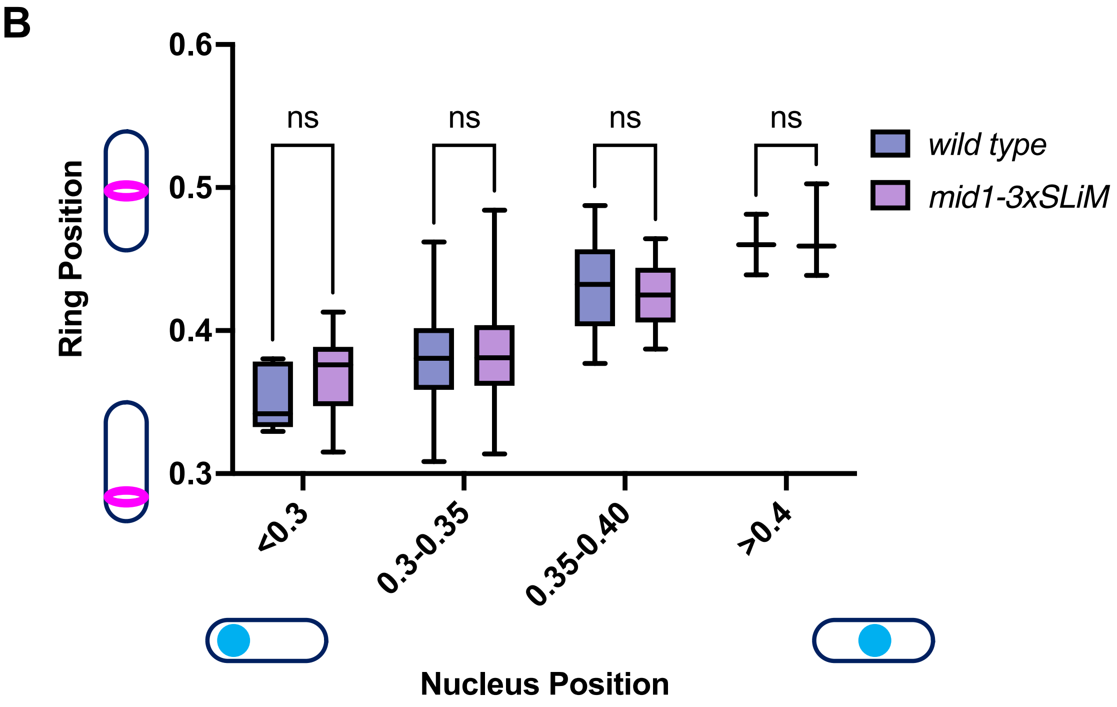
